# A Novel Milli-fluidic Liver Tissue Chip with Continuous Recirculation for Predictive Pharmacokinetics Applications

**DOI:** 10.1101/2023.07.24.550337

**Authors:** Shiny Amala Priya Rajan, Jason Sherfey, Shivam Ohri, Lauren Nichols, J. Tyler Smith, Paarth Parekh, Eugene P. Kadar, Frances Clark, Billy T. George, Lauren Gregory, David Tess, James R. Gosset, Jennifer Liras, Emily Geishecker, R. Scott Obach, Murat Cirit

**Affiliations:** Javelin Biotech Inc., 299 Washington street, Woburn, Massachusetts 01801, United States; Pfizer Global Research and Development, Groton Laboratories, Eastern Point Road, Groton, Connecticut 06340, United States; Pfizer Worldwide Research and Development, 610 Main Street, Cambridge, Massachusetts 02139, United States

**Keywords:** Liver MPS, Primary hepatocyte culture, Pharmacokinetics, induction, in vitro in vivo correlation

## Abstract

A crucial step in lead selection during drug development is accurate estimation and optimization of hepatic clearance using *in vitro* methods. However, current methods are limited by factors such as lack of physiological relevance, short culture/incubation times that are not consistent with drug exposure patterns in patients, use of drug absorbing materials, and evaporation during long-term incubation. To address these technological needs, we developed a novel milli-fluidic human liver tissue chip (LTC) that was designed with continuous media recirculation and optimized for hepatic cultures using human primary hepatocytes. Here, we characterized the LTC using a series of physiologically relevant metrics and test compounds to demonstrate that we could accurately predict the PK of both low and high clearance compounds. The non-biological characterization indicated that the cyclic olefin copolymer (COC)-based LTC exhibited negligible evaporation and minimal non-specific binding of drugs of varying ionic states and lipophilicity. Biologically, the LTC exhibited functional and polarized hepatic culture with sustained metabolic CYP activity for at least 15 days. This long-term culture was then used for drug clearance studies for low and high-clearance compounds for at least 12 days and clearance was estimated for a range of compounds with high in vitro in vivo correlation (IVIVC). We also demonstrated that LTC can be induced by rifampicin, and the culture age had insignificant effect on depletion kinetic and predicted clearance value. Thus, we used advances in bioengineering to develop a novel purpose-built platform with high reproducibility and minimal variability to address unmet needs for PK applications.

## Introduction

Estimating and optimizing the hepatic clearance of small molecule drugs using preclinical pharmacokinetic (PK) testing is a critical step in candidate lead selection. As part of an overall ADME (absorption, distribution, metabolism, and excretion) assessment [1], each drug candidate is assessed to understand the hepatic clearance mechanisms, extent of clearance, and any potential drug-drug interactions (DDI) prior to nominating a lead drug candidate.

Although several cell-based systems, such as suspension hepatocytes and static plated hepatic mono- and co-cultures, are considered current standards for in vitro PK testing for decades, these each possess shortcomings, as detailed by the IQ microphysiological systems (MPS) consortium [2]. For *in vitro* hepatic systems, these unmet needs are described as improved clearance predictions of cytochrome P450 (CYP) and non-CYP metabolized drugs including low clearance compounds, quantification of hepatic clearance mechanisms including intrinsic, uptake, and biliary clearances, and accurate predictions of PK DDI, including induction and time-dependent inhibition.

Suspension hepatocyte cultures have several advantages like convenience and ease of culturing, expressing necessary metabolic enzymes, and allow high cell concentrations. However, they lack physiological relevance, such as efflux transporters expression, and have <6-hour viability, limiting the extent of metabolism detected and effects of longer duration drug exposure. Plated hepatic cultures are more physiologically relevant because they express the relevant transporters and enzymes and form bile canaliculi, which allows biliary clearance assessment. While mono-cultures lose metabolic activity rapidly [3, 4], hepatic co-culture models can extend the metabolic activity, but provide limited flexibility for tissue morphology, and data interpretation can be complicated by the presence of non-human stromal cells [2, 5, 6].

Tissue chips, or MPS, are generally engineered fluidic devices that uses a combination of human cells and biomaterials to better recapitulate the complexity of human physiology [7, 8]. These systems have potential for better assessing PK parameters vs. previous technologies, increasing human physiological relevance with primary human cells, and extending metabolic stability and culture duration. However, common issues that have limited their use in PK studies include single pass flow-through fluidic design (i.e., limited drug exposure to study low clearance drugs), use of drug adsorbing materials for chip fabrication (non-specific binding [NSB] of lipophilic drugs to PDMS), small media volumes (limited kinetic data generation), tissue sizes (limited metabolism for detection), and evaporation during long-term drug incubations [2, 7, 8].

To address these technological needs, we developed a novel milli-fluidic human liver tissue chip (LTC) made of cyclic olefin copolymer (COC) that was designed and optimized for human primary hepatocytes (PHH) cultures maintained under continuous media recirculation. Here, we evaluated this unique system through non-biological characterization (NSB of small molecule drugs and evaporation), biological characterization (tissue morphology and functionality, gene expression, and CYP enzymatic activities), and context-of-use specific characterization (induction, on-chip intrinsic clearance estimation, reproducibility, and in vitro in vivo correlation [IVIVC]). These study findings are also compared to suggestions of the IQ MPS consortium regarding MPS for ADME-related applications [2].

## Materials and Methods

### LTC Culture

Cryopreserved PHHs (BioIVT, Lot WDH, Male,) were seeded as Sandwich Cultured Hepatocyte (SCH) at a density of 215,000 cells/ cm^2^ directly onto the collagen I- and fibronectin- (Corning; Sigma) coated cell chamber, with >90% attachment efficiency. Plated cells were washed with cold culture media (Gibco), overlaid with Matrigel (Corning), and maintained in hepatocyte maintenance media (HMM, Gibco). Next day, 1.7-mL of media was added to the chip after the culture chamber was sealed, and the chips were set to flow at 2mL/h on the controller. LTCs were maintained for long-term culture by replacing 400 μL spent media every 2–3 days and monitored by brightfield microscopy.

### Phenotypic Characterization

Albumin production was quantified in the supernatant using human albumin immunoassays (Meso Scale Discovery) and urea was measured using a colorimetric assay (BioAssay Systems). Concentrations were corrected for number of cells, chip volume, and partial media change volume and frequency to yield daily production rates/million cells.

CYP enzyme activity was assessed with a probe substrate cocktail ( Table S1) metabolized by the major CYP isoforms [9-11]. Hepatic cultures were incubated with the probe cocktail, and metabolism was assessed by metabolite quantification by LC-MS/MS.

Hepatocyte-specific markers, including E-Cadherin (Abcam), F-Actin (Cayman Chemicals), CK-18 (Abcam), MRP-2 (Fisher Scientific), and CYP3A4 (ThermoFisher) were visualized with immunocytochemical staining and fluorescence microscopy. Bile canaliculi were observed with a live staining technique using 5-chloromethylfluorescein diacetate (CMFDA, Invitrogen). Immunofluorescent and brightfield images were taken using an inverted epifluorescence microscope (Zeiss).

### Quantitative real-time PCR (qPCR)

Total RNA was isolated from cells (Invitrogen; PureLink RNA Mini), converted to cDNA (Applied Biosystems; High-Capacity cDNA Reverse Transcription Kit), loaded into a custom Taqman Array plate (ThermoFisher), and analyzed using StepOnePlus Real-Time PCR System (ThermoFisher). Threshold Ct values for each sample were set at 35 for the analysis. RPLO and 18S were selected as house-keeping genes and their expression levels were calculated in parallel with the genes of interest.

### Drug Metabolism Studies

Test compounds were added as cocktails (Table S2) at an initial concentration of 1 µM each, except propranolol at 0.1 µM, during a full medium change, and the final solvent (dimethylsulfoxide-DMSO) concentration never exceeded 0.1% (v/v). Media samples (25–50 µL) were collected from the sampling port at predetermined sampling times (e.g., 1, 4, 24, up to 196 hr). Tissues were then lysed with methanol for intracellular drug quantification. To quantify NSB, drug cocktails of 0.1 - 1 µM concentration were incubated under flow in the chip in the absence of liver tissue and timepoint samples were collected at 24hr. Drug concentrations for both studies were quantified using LC-MS/MS (Table S3 – S7).

### Induction studies

Hepatic cultures in LTC were dosed with 10 µM rifampicin, a known inducer, at day 3 for 72 hr. CYP activity was assessed using a probe substrate cocktail and expression of PK-relevant genes was quantified by qPCR after induction. All inducer-treated tissues were compared to an appropriate DMSO control and inducibility was reported as a fold difference vs. vehicle control. Detailed methodologies (including liver MPS preparation, phenotypic characterization, immunohistochemistry, LC-MS/MS and statistical analysis) are provided in Supplementary Material.

### PK Analysis of Drug Depletion Data

The drug depletion data were analyzed using MATLAB R2019b (MathWorks, Inc., Natick, MA). A one-compartment PK model (model of mono-exponential decay) was used to fit the drug depletion data (Eq. 1):

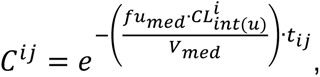

where *C*^*ij*^ is the model prediction for the j^th^ observed concentration regarding the i^th^ chip, sampled at time *t*^*ij*^; *V*_*med*_ is the volume of the medium during the substrate depletion experiment; fu_med_ is the fraction of drug which is unbound in the medium and thus available for metabolism (experimentally determined with equilibrium dialysis); and 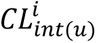 is the unbound intrinsic clearance regarding the i^th^ chip. For NSB correction, the analysis was repeated for on-chip drug study in the absence of the MPS, and the resulting tissue-free value was subtracted. The data from each chip and drug were analyzed independently.

### Prediction of In Vivo Intrinsic Clearance

All related methodological details and the complete procedure (equations) used for in vitro–in vivo extrapolation of clearance is provided in the Supplementary Material.

## Results

### Effect of fluidic configuration on PK studies: Flow-through or recirculating tissue chips

We investigated the theoretical effects of fluidic configurations, recirculation vs flow-through, on drug depletion kinetics in tissue chips. Using quantitative systems pharmacology (QSP)-based algorithms, two hypothetical tissue chips are assumed to have identical hepatic activity and tissue culture chamber dimensions, but one with flow-through configuration and the other with recirculation. After administering 1 µM drug to each configuration, drug depletion kinetics were simulated, and drug profiles plotted for samples collected from media reservoirs (Figure 1). In flow-through configuration, the drug concentration in the media reservoir increases and plateaus within 2 hours. The magnitude of the plateau value is inversely proportional to clearance rate, i.e., the higher the intrinsic clearance rate, the lower the plateau value. In recirculation configuration, the simulated drug profiles were a function of mono-exponential decay. Drug profiles with a range of clearance values were simulated and an arbitrary low threshold value, which is a minimum of 10% parent drug depletion, was assumed to estimate the lowest detection limits in each configuration for comparison. The lowest clearance value in the flow-through system was 5 µL/min/million cells while in recirculation for 8-day drug incubation; the value that results in at least a 10% drug depletion was 0.08 µL/min/million cells. This demonstrates the sensitivity of recirculating tissue chips to observe wide clearance range and its importance to study kinetics of both low and high clearance compounds.

**Figure 1.**
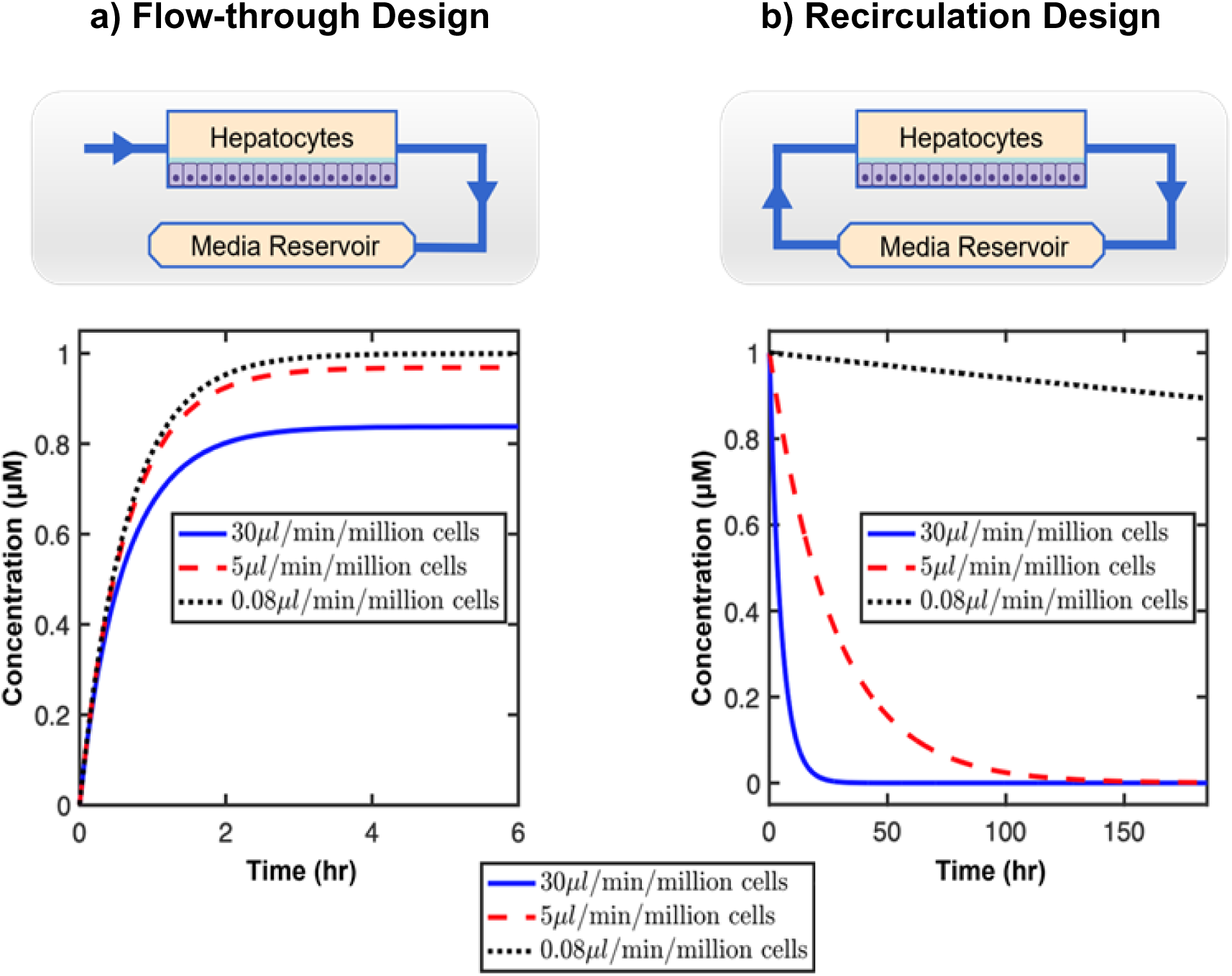
Modeling the effect of recirculation on drug metabolism studies. a) Schematic of flow-through chip design along with the simulated kinetic profile for 1μM parent drug in milli-fluidic LTC without recirculation for three hypothetical compounds spanning a wide range of clearances over 6hrs. b) Schematic of recirculation chip design along with the simulated kinetic profile in LTC with recirculation reveals quantifiable depletion profiles over 8 days

### Recirculating milli-fluidic LTC and LTC controllers

We developed a recirculating milli-fluidic liver tissue chip with an on-board pumping system (Figure 2a). The milli-fluidic design incorporates larger tissue size, media volume, and overall channel dimensions than microfluidic chips. LTC has a 1-cm^2^ cell culture area to accommodate 200,000-250,000 PHH in monolayer for measurable drug metabolism and tissue-based analysis and 1.5–2 mL media volume to allow repeated sampling for drug quantification. Additionally, this design allows multi-scale (media- and tissue-based) PK characterization on a single chip to generate reproducible kinetic and endpoint data. The chip footprint mirrors a standard microtiter plate, making it compatible with standard laboratory equipment.

**Figure 2.**
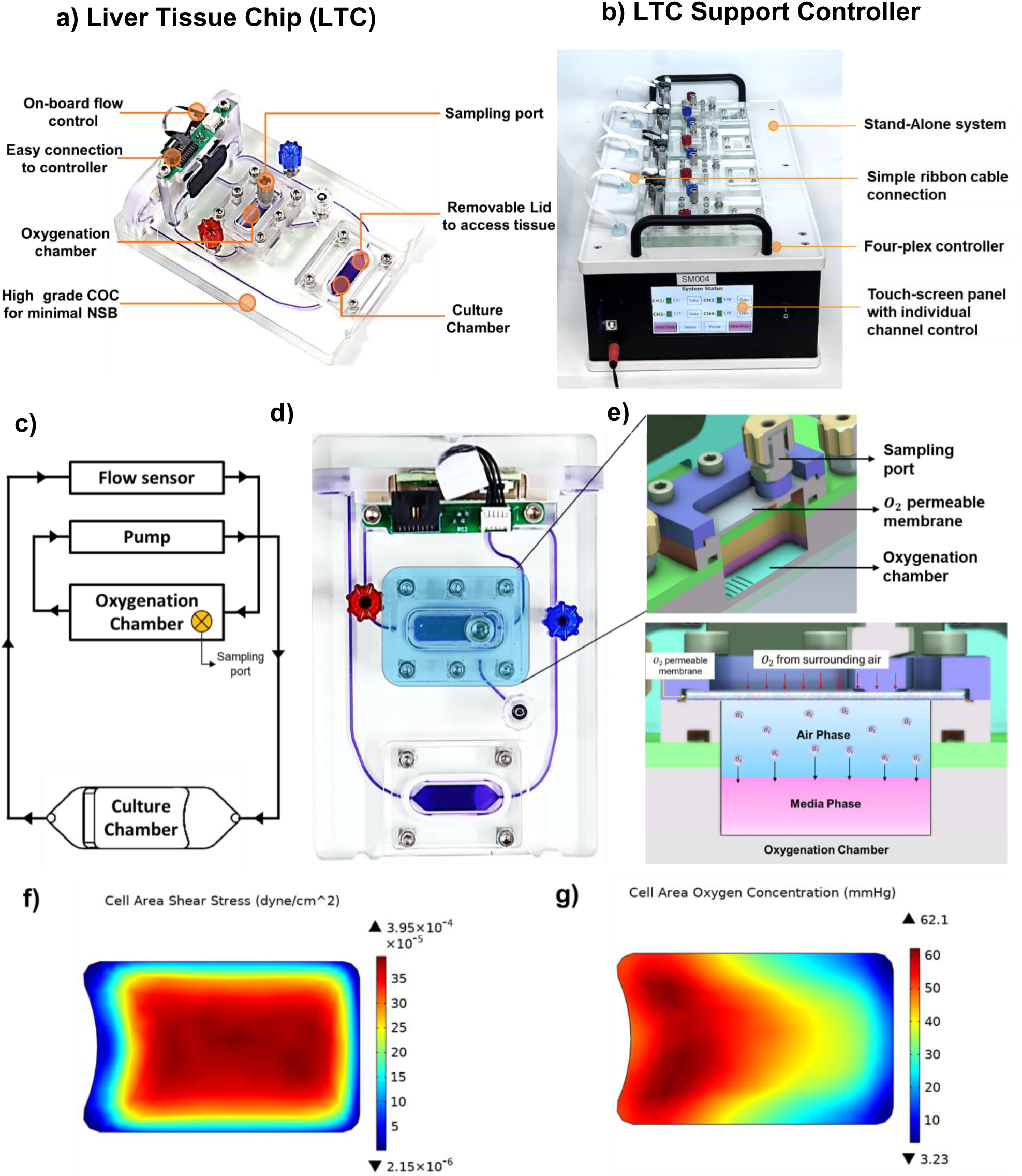
Platform and biomimetic design of the LTC. a) Architecture and key features of the liver tissue chip. b) Features of four-plex controller with LTC connected to flow. c) Schematic of recirculation pathway with d) top view of the LTC. e) Isometric and front sectional view of oxygenation chamber to illustrate media being reoxygenated in the chamber where sampling is carried on during experiment. Computational fluid dynamics (CFD) simulations of LTC culture chamber representing f) a uniform shear profile throughout the tissue achieved by the design parameters in the culture chamber. g) represents the simulation of O2 tension in liver culture chamber that are within the physiologically relevant levels.

LTC has two chambers: a cell culture chamber designed for PHH cultures (mono- or co-cultures) and an oxygenation chamber to re-oxygenate O_2_-depleted medium from the culture chamber that are connected through closed-loop channels and on-board pump system allowing media recirculation (Figure 2b, c, d). Here, the oxygenation chamber allows passive gas exchange between ambient air and air in the oxygenation chamber through an O_2_-permeable silicone membrane. As such, there is no contact between the membrane and recirculating media (Figure 2e), thus avoiding silicone-related NSB of drugs. LTC is designed to mitigate bubble formation issue, which is a challenge in microfluidic systems that has detrimental effects on cells [12-14]. Additionally, the recirculating medium can be accessed with a standard single-channel pipette via the sampling port located on the lid of the oxygenation chamber allowing collection of multiple time points samples, e.g., seven time-points of 50 µL sample volume. The culture chamber is designed with a removable lid that allows direct access to the cell compartment during seeding, longitudinal experiments, and endpoint assays.

The media recirculation was driven by an on-board piezoelectric diaphragm pump eliminating the need for external pumps, reservoirs, and tubing connections. The pump is actuated by an on-board flow sensor that measures the flowrate in real-time and is actively controlled by a four-plex controller which is connected via a ribbon cable (Figure 2b). The controller regulates the recirculation with a continuous feedback loop between the pump and flow sensor to maintain constant and accurate flow. Each chip is controlled independently, and the flow rates can be set with the touch-screen panel. This is a stand-alone and user-friendly platform that does not require any additional external support or computer connection and up to four controllers were used in an incubator.

### Computational investigation of optimal recirculation flowrate in LTC

High oxygen consumption and low shear tolerance are two essential characteristics of primary hepatocyte cultures [15-18]. In LTC, the culture chamber is designed to provide uniform shear stress (Figure 2f) by continuous perfusion of the hepatic culture at appropriate flowrate. As the O_2_ in media entering the cell chamber is consumed by cells, the oxygen-depleted media leaving the chamber gets reoxygenated in the oxygenation chamber through gas exchange. These phenomena were modeled with computational fluid dynamics (CFD) simulations to determine optimal flowrate to bring sufficient O_2_ to the culture, while maintaining low shear (<0.005 dyne/cm^2^). The calculated O_2_ tension on the hepatic tissue is similar to physiologically relevant O_2_ concentrations observed in the human liver (45–50 mmHg in periportal zone 1 and 15–20 mmHg in perivenous zone 3) [19, 20]. CFD-based O_2_ concentration estimates for LTC were 50–60 mmHg at the inlet and 15–25 mmHg at the outlet (Figure 2g). The experimental validation of on-chip O_2_ concentration calculated by CFD will be valuable in future research. Through simulations and empirical experiments, 2 mL/h flow rate was determined to achieve sufficient oxygenation, appropriate nutrient delivery to the hepatic tissue, efficient removal of cellular waste, optimal shear stress and uniform drug distribution in LTC. In addition, the features in chamber inlet minimize the shear from high flow rates to hepatocyte tolerable shear range, that helps to maintain the improved morphology and metabolic activity during long-term culture.

### Non-biological characterization of LTC

In PDMS microfluidic chips, NSB drastically increases with increasing lipophilicity [18], and the absorption increases with the hydrophobicity and topological polar surface area of the small molecule [21, 22]. In contrast, COC is a relatively hydrophilic material vs. PDMS with a higher surface energy. To validate minimal NSB in LTC, small molecule compounds at 0.1 - 1-µM initial concentrations, were recirculated on fully assembled tissue chips without hepatic tissue for 24 h; samples were collected at 0 and 24 h and quantified by LC-MS/MS. Twenty-one compounds of varying ionic state and lipophilicity (logP) ranging from –1.04 to 6.08 were tested and near complete recovery of neutral and basic compounds was achieved, while the high logP acidic drugs (bosentan, pitavastatin, fluvastatin, and repaglinide) was recovered >60% (Figure 3).

**Figure 3.**
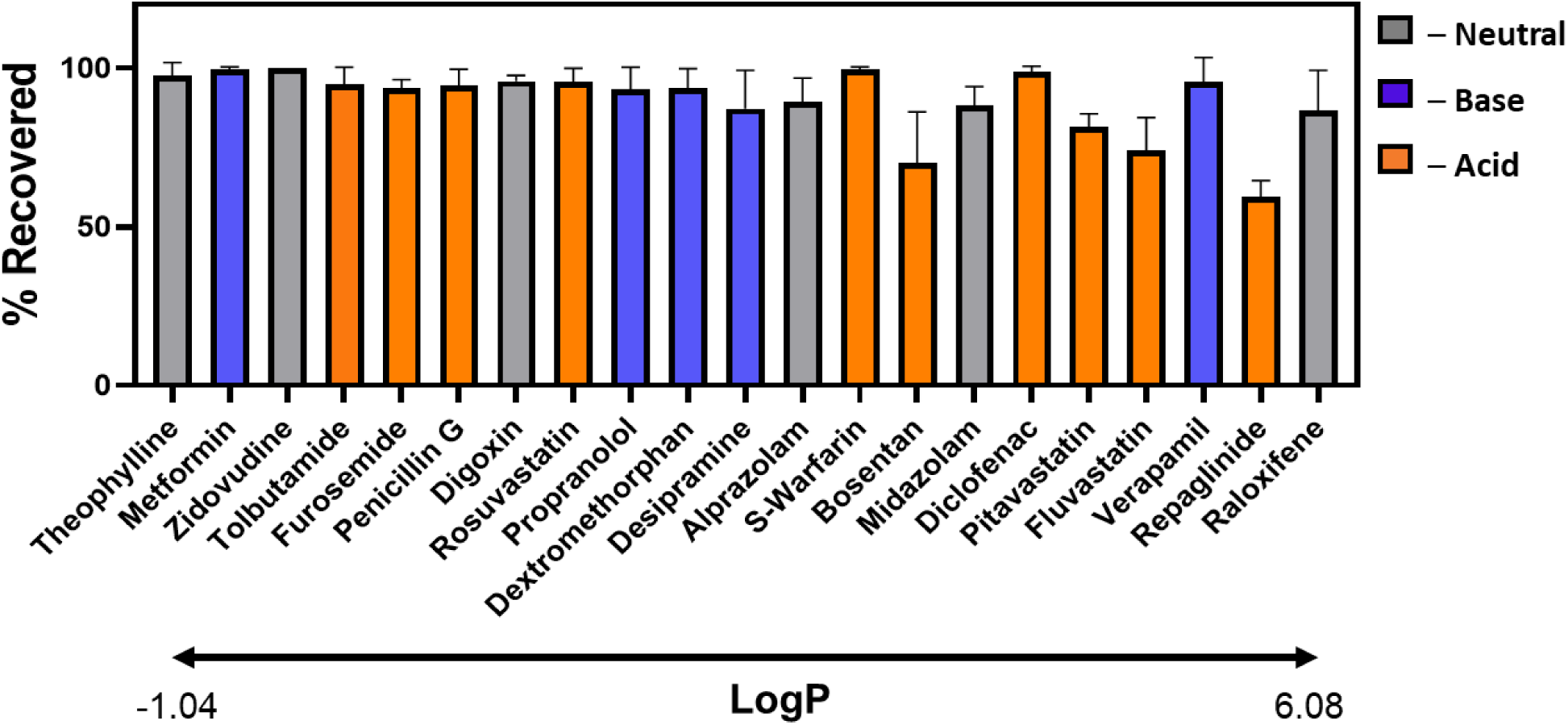
Non-Specific binding (NSB) characterization of Liver Tissue Chip. NSB of various small molecules were determined by incubating 0.1-1μM drug concentration for 24hr under recirculating flow without hepatic tissue. 17 out of 21 drugs were recovered over 85% of their initial concentration.

Evaporation can be a concern for drug studies, usually in micro-bioreactor-type tissue chips in which the media is exposed to ambient air [23-25]. In such systems, continuous perfusion may contribute to higher evaporation which becomes critical for long-term drug incubations, especially when the evaporation rate are in similar order of magnitude to clearance rates [26]. Due to COC’s material property, the evaporative media losses evaluated over a week with no media replenishment were negligible <0.4% per day or <2.8% over a week (Figure S1).

### Biological characterization of LTC

To demonstrate metabolically active long-term hepatic cultures and polarized morphology, which are essential for PK applications, PHH were seeded as a SCH with collagen I and fibronectin as underlay and Matrigel® GFR as overlay to allow polarized hepatocytes for maintenance of differentiated morphology, stabilized metabolic activity, and bile canaliculi formation. After a day, the culture was maintained under continuous recirculation (2 mL/h) for the rest of the experiment and performed partial media change every 2-3 days to allow media conditioning [27-29]. Live imaging revealed that the differentiated morphology of characteristic cobblestone pattern maintained for at least 15 days (Figure 4a). LTC was live stained for formation of bile canaliculi using CMFDA (Figure 4b), which was observed at the junction of cells confirming polarization. IHC staining showed canalicular membrane expressing multidrug resistance-associated protein 2 (MRP2), an apical bile acid transporter. The tissue stained positive in cytoplasm for CYP3A4, and epithelial cytoskeleton marker, CK18, which is highly concentrated in hepatocytes [30, 31] and E-cadherin, an epithelial marker expressed in the adjacent cell boundaries of differentiated hepatocytes as a cobblestone pattern. 594-Phalloid stained the high-density F-actin filaments that are arranged in the apical side of hepatocytes and localized in the bile canaliculi bordering hepatocytes.

**Figure 4.**
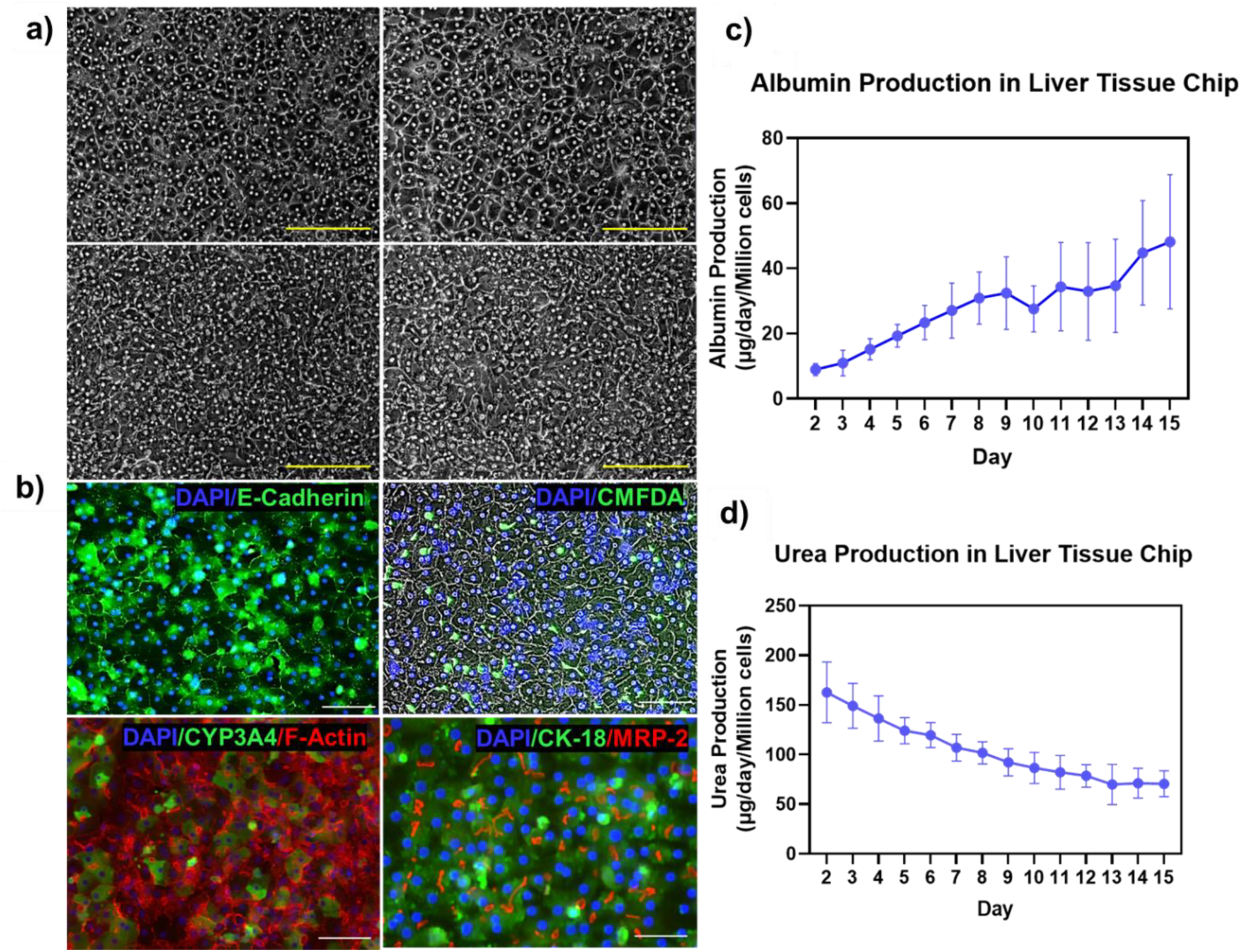
Biological characterization of LTC with morphology imaging and biochemical assays. a) Long-term hepatocyte morphology imaged using bright field microscope on day 3, 7, 11 and 15 showing phenotypic cobblestone morphology. b) Immunocytochemistry staining on day 7 showing presence of E-cadherin, F-Actin, CYP3A4, CK-18, MRP-2 and bile canaliculi using CMFDA staining. c) Albumin and d) urea levels were assessed in our liver chip system daily on days 2–15.

Albumin and urea production rates, which are commonly used to evaluate *in vitro* functionality of hepatocyte cultures, were measured throughout the 15-day culture period (Figure 4c, d). The hepatic culture in LTC has comparable albumin and urea production to estimated human liver production (37-105 μg/day/million cells for albumin production and 56-159 μg/day/million for urea production) [7]. The initial increase in albumin levels can be attributed to gradual remodeling of the ECM in the tissue microenvironment stimulated by recirculation of the conditioned media. While the albumin production rate steadily increased, the urea production rate decreased slowly over 15 days. Gene expression studies (Figure S2) also showed a progressive increase in albumin mRNA levels (>2.5-fold) in the 15-day culture, following similar trends as albumin production.

To demonstrate the biological reproducibility of LTC, we quantified the coefficient of variance (CV%) for albumin and urea levels in four independent experiments on day 3, 8 and 10 (Figure S3a, b). Compared to current MPS models, albumin and urea data shows < 28.5% variability across chips and <29.5% between experiments and operator, leading to a robust and reproduceable platform. (Figure S3c)

The metabolic activity of hepatocyte culture was quantified by assessing CYP isoform-specific metabolite formation at days 3, 8, and 13 using CYP-probe cocktail (Table S1). CYP1A2, CYP2C9, CYP2C19, CYP2D6, and CYP3A4 retained or increased activity for up to 13 days of culture (Figure 5a). The quantified drug metabolizing Phase 1 enzymes (*CYP Isoforms, CES1 and CES2*), Phase 2 (*UGTs, GST*) and transporter (*SLCs, ABCs*) genes using RT-PCR maintained their expression levels up to 15 days, with varying degrees of fold-change (Figure 5b).

**Figure 5.**
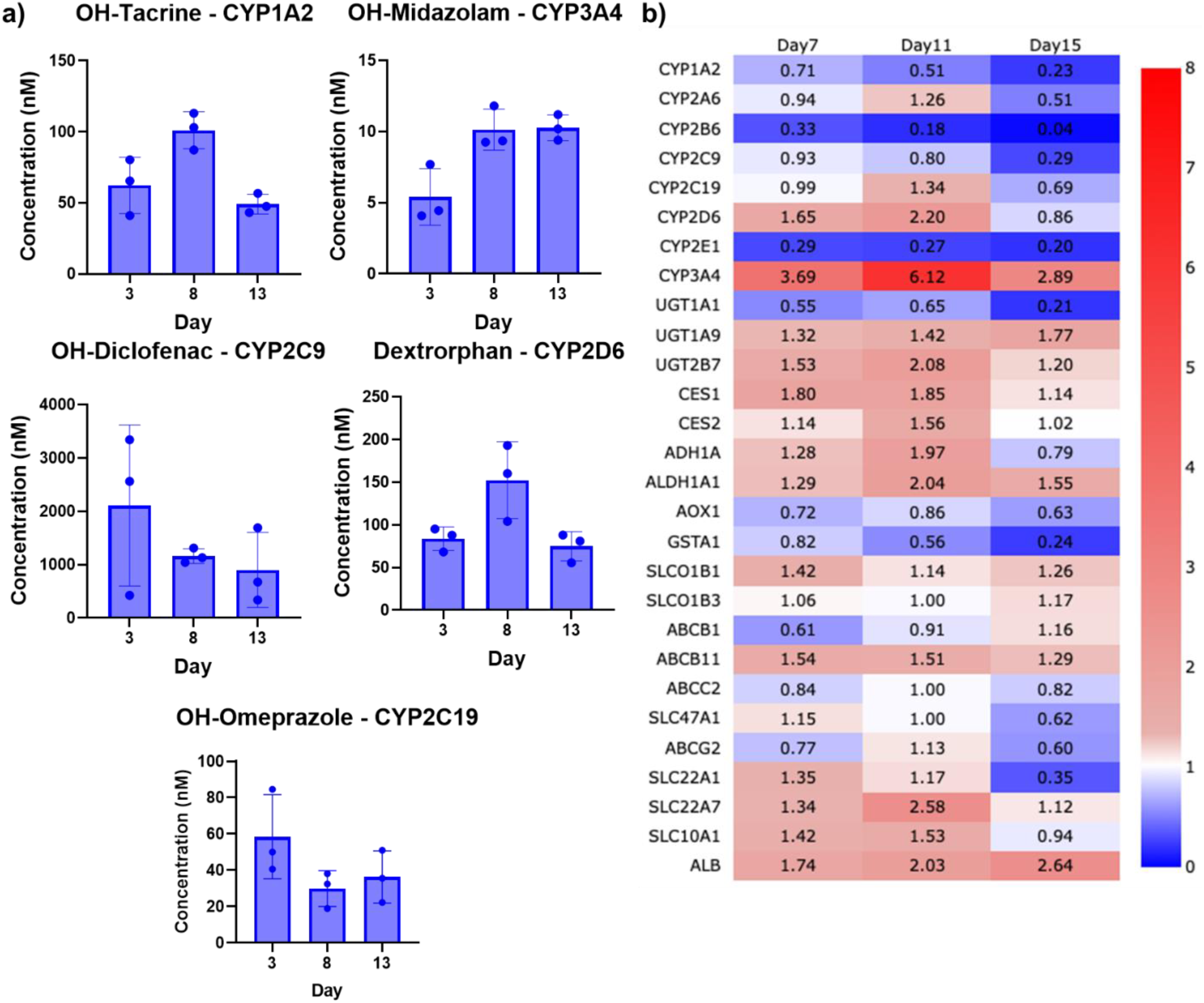
Activity and gene expression levels of cytochrome P450 isoforms in liver tissue chip. a) The relative activity levels and stability of CYP1A2, CYP3A4, CYP2C9, CYP2D6 and CYP2C19 CYP isoforms were measured by cocktail CYP probe substrate analytical assay. b) Fold change expression as measured by TaqMan RT-PCR on days 7,11, and 15 relative to day3 for the PK-relevant genes

Here, we demonstrated that LTC’s PHH culture maintained their polarized morphology and functional markers comparable to *in vivo* for at least 15 days, while the long-term culture retained metabolic activity measured as both drug-metabolizing enzyme levels and gene expression profiles of PK-relevant genes.

### Effect of culture age and drug incubation window on intrinsic clearance estimation in LTC

Intrinsic clearance values were evaluated by incubating cocktails of four high-clearance compounds (midazolam, propranolol, diclofenac, and dextromethorphan) on days 3, 5, or 7 of culture for 72 h and two low-clearance compounds (alprazolam and tolbutamide) on days 3 and 7 for 120 h. High clearance drugs were significantly depleted over 72 h, whereas low clearance drugs took up to 120 h for 20% and 50% parent drug depletion, respectively (Figure 6). The intrinsic metabolic clearance (*CL*_*int*(*u*)_) values for each drug and drug incubation window (Table 1) shows high reproducibility with average CV% of 18% and no statistically significant differences in variance or AUC for any compounds. However, ***CL***_***int***_for diclofenac (p=0.052) and propranolol (p=0.11) slightly decreased across culture windows which may be related to changes in CYP2C9 activity levels and *gene* expression over time (Figure 5a, b) and other related metabolizing enzymes. Overall, LTC produces very consistent drug depletion profiles and ***CL***_***int***_ estimates with minimal chip-to-chip and day-to-day variability. Thus, chips can be potentially used for repeated drug studies for low and high-clearance compounds at least for 12 days of culture and has flexibility regarding the culture window used for drug depletion studies of high and low cleared drugs.

**Figure 6.**
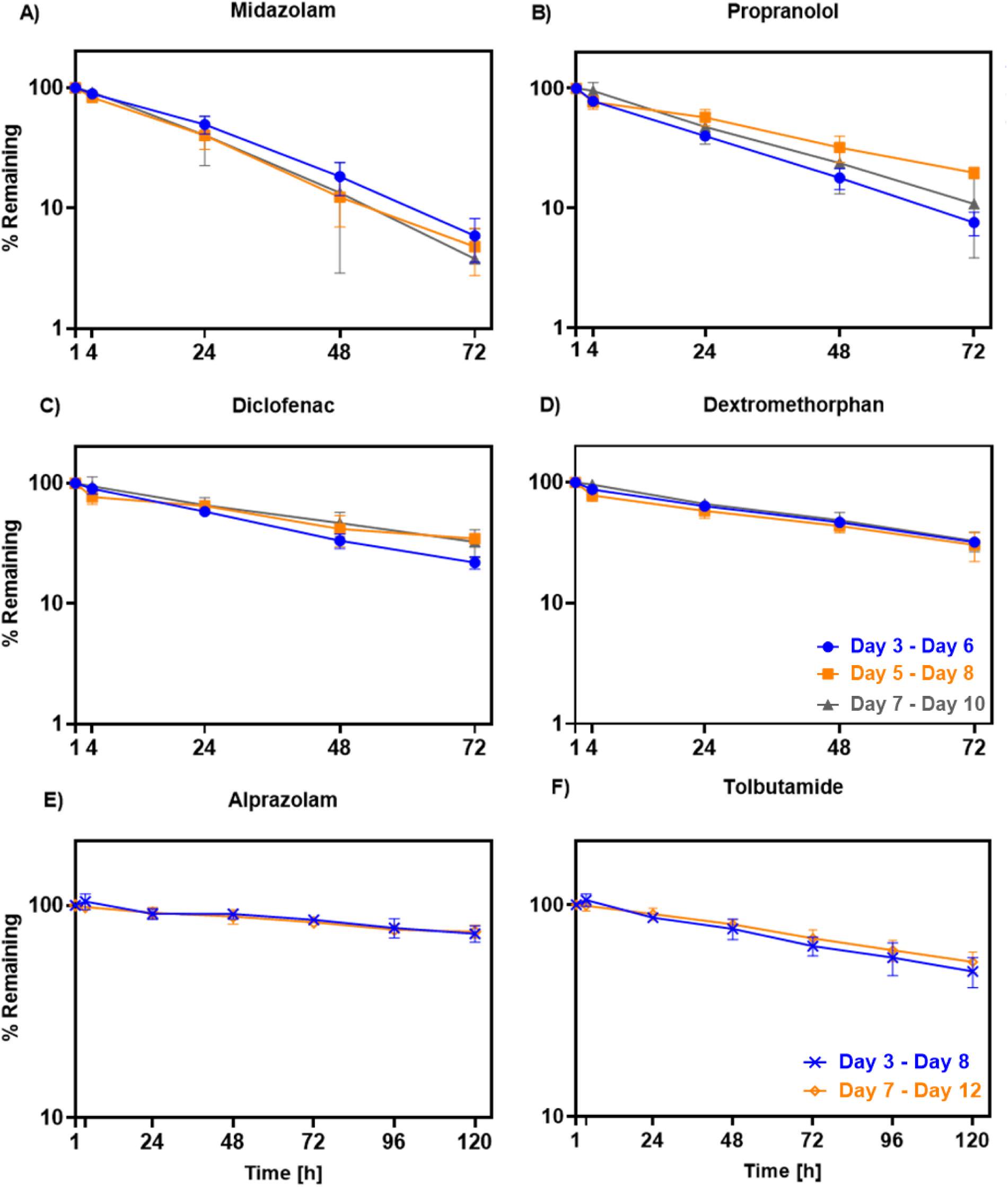
Effects of culture age and drug incubation on intrinsic clearance studies in liver tissue chips. Both high clearance drug [a) Midazolam, b) Propranolol, c) Diclofenac and d) Dextromethorphan] and low clearance drugs [e) Alprazolam and f) Tolbutamide] were evaluated on LTC at different incubation window for 3 and 5 days of incubation period respectively.

**Table 1.** Intrinsic metabolic clearances (*CL*_*int*_ [μL/min/million cells]) calculated from LTC with different incubation windows.

| **Drug** | **Incubation Window** | **n** | **Mean ± SD**<br>[μL/min/million<br>cells] | **CV** |
| --- | --- | --- | --- | --- |
| **Alprazolam** | Day 3 - Day 8 | 4 | 0.54 ± 0.23 | 36% |
| **Alprazolam** | Day 7 - Day 12 | 5 | 0.51 ± 0.10 | 18% |
| **Dextromethorphan** | Day 3 - Day 6 | 3 | 2.86 ± 0.07 | 2.3% |
| **Dextromethorphan** | Day 5 - Day 8 | 3 | 2.87 ± 0.68 | 21% |
| **Dextromethorphan** | Day 7 - Day 10 | 4 | 2.91 ± 0.57 | 18% |
| **Diclofenac** | Day 3 - Day 6 | 3 | 81.8 ± 7.50 | 8.8% |
| **Diclofenac** | Day 5 - Day 8 | 3 | 54.0 ± 3.80 | 6.9% |
| **Diclofenac** | Day 7 - Day 10 | 4 | 59.7 ± 17 | 26% |
| **Midazolam** | Day 3 - Day 6 | 3 | 12.0 ± 2.10 | 16% |
| **Midazolam** | Day 5 - Day 8 | 3 | 13.2 ± 2.20 | 16% |
| **Midazolam** | Day 7 - Day 10 | 4 | 14.8 ± 4.80 | 29% |
| **Propranolol** | Day 3 - Day 6 | 3 | 5.34 ± 0.53 | 9.4% |
| **Propranolol** | Day 5 - Day 8 | 3 | 3.26 ± 0.51 | 15% |
| **Propranolol** | Day 7 - Day 10 | 4 | 4.99 ± 1.80 | 31% |
| **Tolbutamide** | Day 3 - Day 8 | 4 | 1.55 ± 0.48 | 28% |
| **Tolbutamide** | Day 7 - Day 12 | 5 | 1.30 ± 0.21 | 15% |

### Effect of Rifampicin induction on LTC

LTC were dosed with 10 mM rifampicin on day 3 and incubated for 3 days. Induction increased metabolic activity of CYP2C9, CYP3A4, and CYP2C19, while CYP2D6 activity was unchanged (Figure 7a). Additionally, mRNA analysis showed differential effects of induction on various Phase 1 drug-metabolizing enzymes, conjugative UGT enzymes and efflux Transporters (ABCs) (Figure 7b).

**Figure 7.**
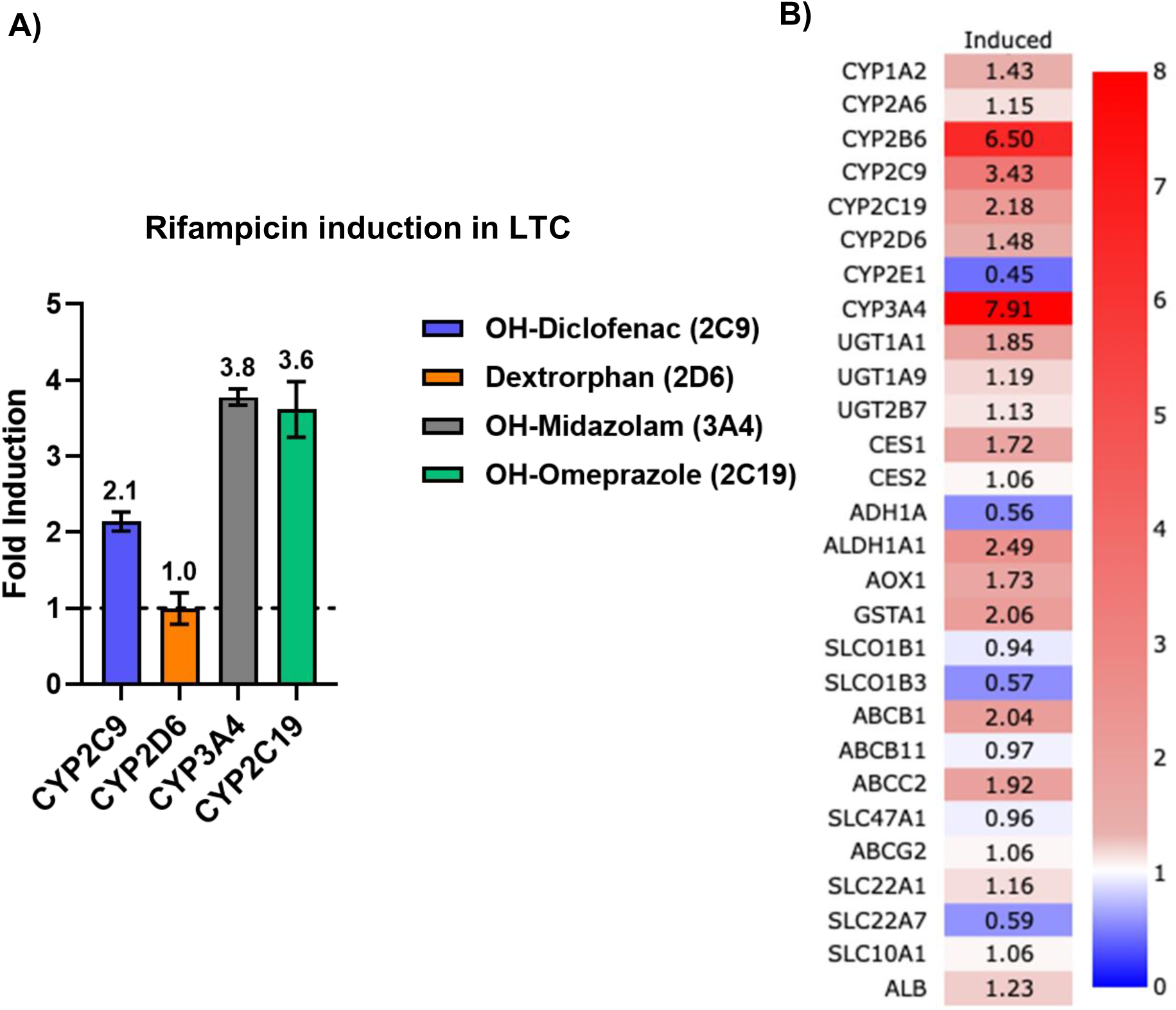
Effects of Rifampicin induction on CYP enzyme activity and mRNA expression of PK relevant gene. a) cocktail CYP probe substrate analytical assay and b) mRNA expression levels as measured by TaqMan RT-PCR. The fold change expression levels of the induced group activity levels are relative to the DMSO vehicle control group.

### Estimation of intrinsic clearance values of small molecule drugs and IVIVC

A range of compounds with varying clearance values, physicochemical properties, and metabolizing enzyme-specificity were assayed in LTC to estimate on-chip clearance values (Table S8). Observed intrinsic clearances spanned three orders of magnitude from 0.15 μL/min/million cells for rosuvastatin to 115 μL/min/million cells for repaglinide (Figure 8). Diclofenac and S-warfarin, both metabolized by CYP2C9, showed intrinsic clearances of 81.8 μl/min/million cells and 0.82 μl/min/million cells, and raloxifene and zidovudine, both metabolized by UGTs, showed intrinsic clearances of 70.6 μl/min/million cells and 1.94 μl/min/million cells, showing differential clearance values for both CYP and UGT substrates. For all drugs, the standard deviation across chips was small compared to the mean (13% median CV; 16% mean CV, excluding rosuvastatin, which had 440% CV and the lowest CL_int_ value).

**Figure 8.**
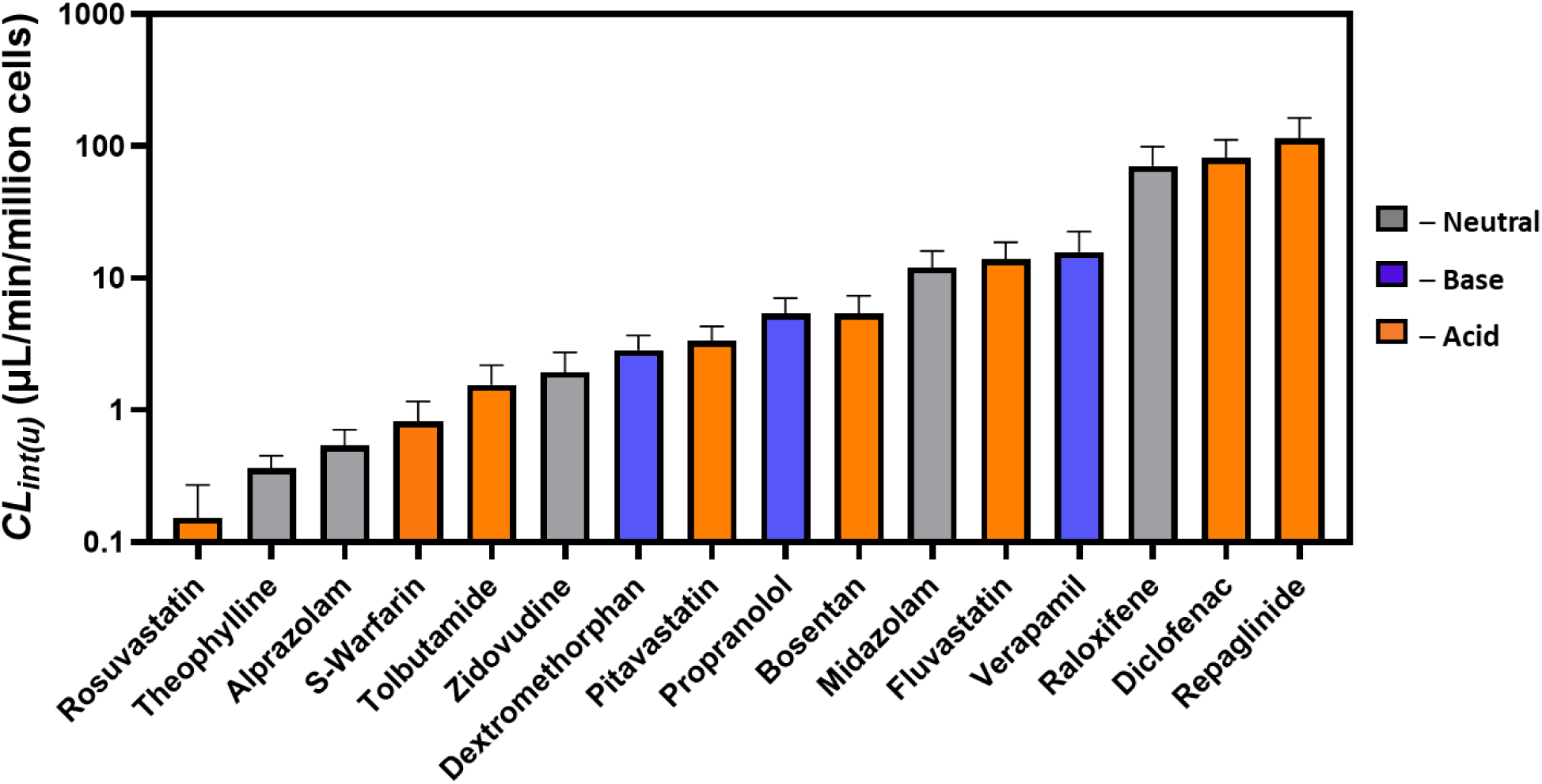
Unbound intrinsic clearance (*CL*_*int*(*u*)_) for 16 compounds estimated from LTC. Error bars show standard deviations across chips. The bar color indicates whether the compound is an acid (orange), base (blue), or neutral (gray)

To evaluate the clinical relevance of the clearance values, *in vitro* intrinsic clearance values from LTC were scaled to predicted human clearance values using human hepatocellularity and the parallel tube model (PT, Table S9). Figure 9a shows the IVIVC of intrinsic clearance of ten drugs for which clinical intrinsic clearance values can be estimated. IVIVC of these drugs showed average absolute fold error (AAFE) of 2.7, and 40% of the predictions were within 2-fold of clinically derived values and were particularly well-predicted for drugs with low intrinsic clearance.

**Figure 9.**
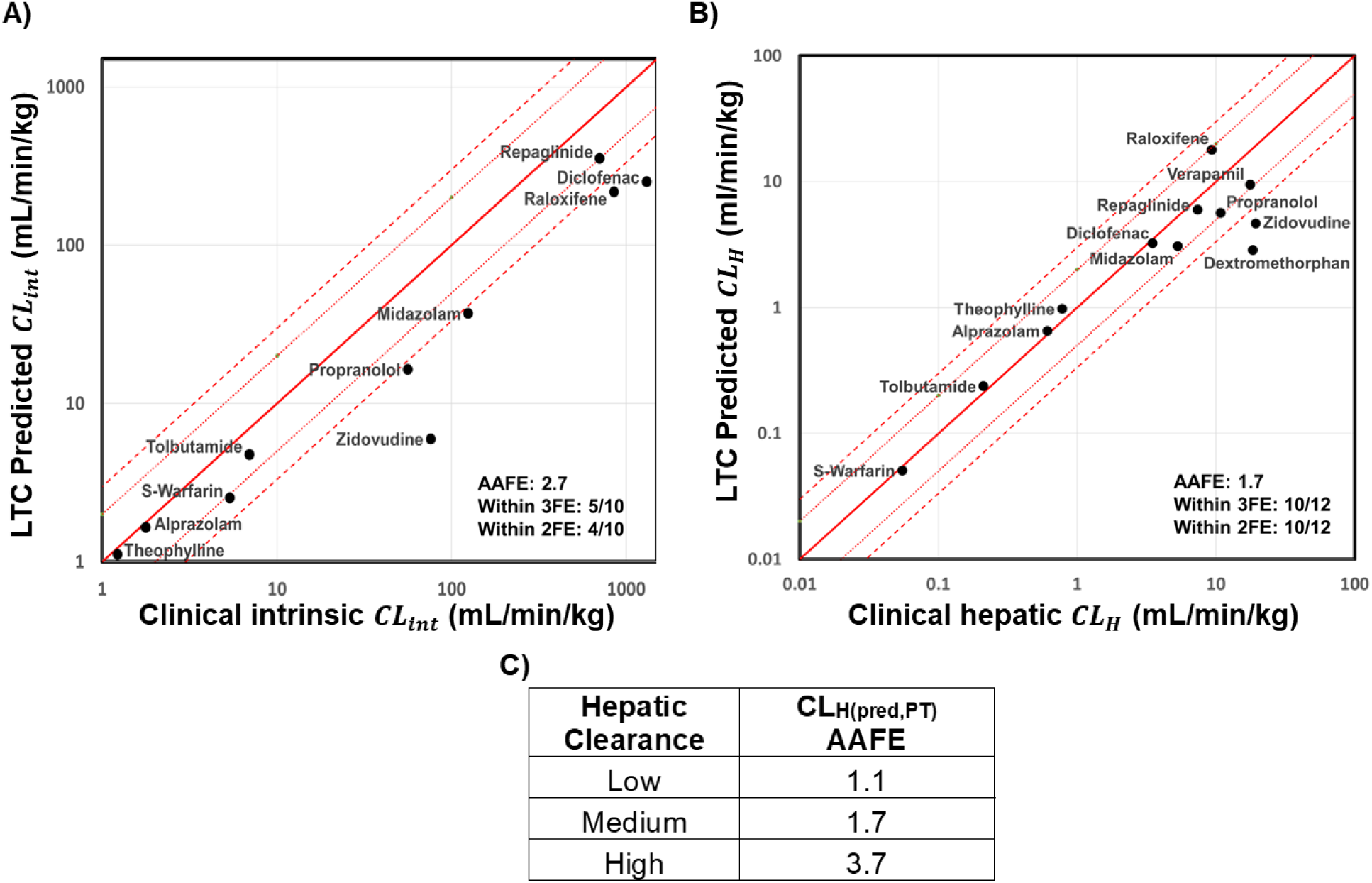
In vitro in vivo correlation (IVIVC) of LTC predicted clinical clearance estimates to clinically observed values for A) intrinsic metabolic clearance (*CL*_*int*_) and B) total hepatic clearance (*CL*_*H*_). LTC estimates of human *CL*_*int*_were obtained by scaling the observed in vitro LTC *CL*_*int*_ according to human hepatocellularity. LTC estimates of human *CL*_*H*_ were obtained by further application of the parallel tube model. C) The table shows average absolute fold errors calculated across drugs with different clearances shown in 9B. The *in vivo* human hepatic clearance for drugs are classified as low clearance (*CL*_*H*(*obs*)_< 5 ml/min/kg), medium clearance (5 < *CL*_*H*(*obs*)_<15 ml/min/kg), and high clearance (*CL*_*H*(*obs*)_> 15 ml/min/kg).

After applying the PT model, which accounts for blood flow and protein binding, the in vivo hepatic clearance (CL_H_) due to metabolism is better predicted (Figure 9b). For the 12 drugs cleared primarily hepatically, the LTC-predicted in vivo hepatic clearance values with AAFE of 1.7, and 83% of the predictions were within 2-fold of clinical values and comparable results were obtained using the well-stirred model (1.8 AAFE, see Table S8). The in vivo hepatic clearance was best estimated for drugs with low clearance (S-warfarin, tolbutamide, alprazolam, and theophylline). AAFE for low, medium, and high clearance drugs were 1.1, 1.7, and 3.7, respectively (Figure 9c). Most high-cleared compounds were predicted within 2-fold of clinical values except zidovudine and dextromethorphan. Additionally, for few compounds with detectable NSB (>10% depletion over 24 h), a computational correction was applied to account for drug depletion due to NSB, which had minimal impact on the estimate: with 16% on average shift with NSB correction. These results demonstrate that the LTC generates highly reproducible drug depletion kinetics and accurate estimates of clinical clearance values for diverse set of compounds.

## Discussion

The tissue chip field has been emerging for the past decade and is expected to gain more momentum and traction after the 2022 FDA Modernization Act. With recent advances in bioengineering, tissue chips can be purpose-built as new approach methods (NAMs) for certain context of use (CoU) applications to improve performance of current preclinical *in vitro* and animal models. Qualification and validation are critical aspects of the development and implementation of new tools or methods in drug development.

Validation of NAMs, such as tissue chips, ensures that the technology and their intended (CoU) applications are reliable and provide accurate data for regulatory decision-making. FDA’s Drug Development Tools (DDT) and Innovative Science and Technology Approaches for New Drugs (ISTAND) programs provide guidelines about the needs for validation of NAMs. A qualified DDT can be relied upon to have a specific interpretation and application in drug development and regulatory review within its stated CoU. Once qualified, DDTs will be available to use in drug development programs for the qualified CoU.

While many applications of tissue chips have been developed, these mainly focus on healthy and disease tissue models in toxicology and efficacy testing. However, drug metabolism applications to study metabolite profiling, prediction of PK parameters, and DDI potential, has been limited. The unmet needs have been discussed and summarized by the IQ MPS consortium [2]. Briefly, i) culture media recirculation to allow adequate metabolic turnover of test compounds; ii) tissue chip material that exhibits low NSB; iii) sufficient system and tissue volume for media-based multiple sampling; iv) sufficient tissue mass to observe metabolism and for end-point tissue-based analysis; v) low evaporation for long-term incubation and continuous sampling; and vi) sufficient throughput to run on-chip PK studies. Here, we used advances in bioengineering to address these unmet needs for PK applications by designing and developing a novel purpose-built milli-fluidic COC-based LTC with continuous recirculation.

The LTC is completely PDMS-free and fabricated with >90% thermoplastics, of which 82% is high-grade COC with superior mechanical and chemical properties [32]. This COC-based chip construction reduces the NSB of small molecules; however, some compounds with high lipophilicity may retain some degree of NSB. In those cases, to improve the accuracy of the clearance predictions, it is important to evaluate the test compounds for NSB in tissue chip and throughout the analytical workstream, including pipette tips, sample vessels, and LC-MS/MS. The superior moisture barrier property of COC and the chip architecture significantly reduces media evaporative losses, which can be an issue for long-term drug incubation with well-plates and in micro-bioreactor-type tissue chips [24]. LTC also allows for additional design capabilities such as removable culture chamber lid and incorporation of auxiliary features such as on-board pumps, sensors, and sampling ports. The removable lid allows researchers to directly access the culture chamber during seeding, throughout the experiment for non-sacrificial tissue-based assays, e.g., CYP activity assays, and for endpoint assays, e.g., transcriptomics or immunohistochemistry assays. The sampling ports are designed to collect accurate media volumes from the oxygenation chamber using standard pipettes to collect multiple samples throughout studies for kinetic data generation. For PK studies in this manuscript, we routinely ran high and low clearance drugs for 3- and 5-day incubations, respectively, and demonstrated up to 8-day drug incubation (Figure S4). In each study, we collected multiple 25–50 mL samples from a single chip without replacing the medium that has the advantages for data quality over collecting each timepoint from different wells, which increases well-to-well variability due to cellular variability in each well, e.g., cell numbers.

Demonstration of reproducibility of a NAM is crucial for wide-spread adoption of LTC systems. Here, LTC reproducibility was assessed for three main criteria: i) non-biological functions of tissue chips (flow rate and NSB); ii) Biological functions (morphology, albumin & urea production, and gene expression profiles of hepatic tissue); and iii) CoU-specific studies (CYP-activity, induction and intrinsic clearance estimation of small molecule drugs). For all three criteria, LTC demonstrated high reproducibility and low experiment variability. It is important to note that while biological characterization may show expected changes over the time of tissue function *in vitro*, for example increased albumin levels and decreased urea levels over 15 days, CoU-specific reproducibility assessment showed no statistically significant change in clearance estimation starting drug incubation at different culture ages. Therefore, it is important to perform reproducibility experiments not only for biological function of NAM, but also for intended CoU applications. For future work, assessment of donor variability is critical to demonstrate the expected donor-to-donor variability and establish pooled donors for general or subpopulations. Additionally, assessment of lab-to-lab variability is necessary to accelerate the adoption of NAMs.

Another important aspect of assessing tissue chip performance is to evaluate its predictive power for the intended CoU. Here, we demonstrated that on-chip *CL*_*int*_values of small molecules can be scaled to clinical clearance values and be compared directly using quantitative IVIVC plots (correlating clinically observed values to predicted values from LTC experiments). The predictive power of different experimental systems that are used to evaluate intrinsic clearance values can also be compared using IVIVC methodology. The value of highly correlative NAM may increase confidence in decision-making and, hence, reduce the need for additional experimentation, such as animal studies. Furthermore, the intrinsic clearance parameters can be used with PBPK and QSP models to estimate first-in-human dosing.

While we focused on PK CoU of small molecule drugs with LTC using PHH in SCH format, LTC can accommodate various hepatic cytoarchitectures including 3D mono/ co-cultures with non-parenchymal cell types (e.g., Kupffer cells, stellate cells, and liver sinusoidal endothelial cells) and applied to various CoUs. These CoU applications can be within the needs of PK departments, e.g., inhibition and induction studies with low clearance drugs or PK studies with diseased liver phenotypes, and different needs in pharmacology (e.g., infectious liver disease) and toxicology (e.g., drug-induced liver injury). As a proof-of-concept, we also demonstrated the inducibility of LTC with rifampicin and its effect on PK-relevant genes. These studies provide essential preliminary data to make informed decisions about further advancement in testing of novel compounds. LTC may also be used to test other modalities and drug delivery systems, such as large molecules, oligonucleotides & LNPs, and gene therapies & AAVs. The human specificity of the hepatic tissue may be valuable for modalities against human-specific targets. We maintain that the complexity of the hepatic cytoarchitecture should be carefully determined based on the needs of the intended CoU applications and be diligently assessed for applicability and reproducibility.

This work is also an important building block for multi-tissue chips with integrated system pharmacology [33,34]. It is crucial to have metabolically active tissue for PK applications using multi-tissue chips, which can include combinations of PK-relevant tissue models, e.g., gut, liver, kidney, blood brain barrier MPSs. Such multi-tissue systems could potentially be used to evaluate multiple ADME properties and biotransformation. Moreover, the design of the LTC allows incorporation of sensors in the fluidic path. While flow sensors are already incorporated in LTC and controller design, in future designs, additional sensor technologies, e.g., O_2_ and pH sensors, can be added to the system to generate real-time chip data.

## Conclusion

Here, we have described the design and construction of a milli-fluidic liver chip and focused on several aspects important to the use of the platform in evaluation of drug disposition (e.g. NSB, long duration incubations, suitable volumes for sampling, etc) [2]. The system successfully demonstrated long-term maintenance of hepatocyte physiology and ability to project human drug clearance, especially for difficult-to-measure low intrinsic clearance drugs with high correlations. This system offers an important advancement to improve drug research while simultaneously reducing the need for experiments in laboratory animals and will serve as a springboard to the construction of integrated multi-organ systems that will recapitulate the overall disposition of drugs in humans.

## Acknowledgments

The authors thank Dr. Donald Tweedie for providing invaluable guidance and discussions during the manuscript review. We are also grateful to Anna Kopec, Manthena Varma, and Christine Orozco for comments and discussions, and to Ramona Morales and Alexis Runstadler for laboratory support.

## Author Contributions

S.R., J.S., and M.C. wrote the manuscript; M.C., S.R., J.S., E.G., R.S.O., D.T., J.R.G., J.L., designed the research and computational modelling; S.R., S.O., and L.N., performed the research; J.S., J.T.S, and P.P., analyzed the data and performed computational modelling; E.P.K., F.C., B.T.G., LG performed all the bioanalytical work. All authors discussed the results and reviewed the manuscript.

