## Supplementary materials for "A Novel Milli-fluidic Liver Tissue Chip with Continuous Recirculation for Predictive Pharmacokinetics Applications"

Supplementary material for the manuscript “A Novel Milli-fluidic Liver Tissue Chip with Continuous Recirculation for Predictive Pharmacokinetics Applications”

Shiny Amala Priya Rajan^1^*, Jason Sherfey^1^*, Shivam Ohri^1^, Lauren Nichols^1^, J. Tyler Smith^1^, Paarth Parekh^1^, Eugene P. Kadar^2^, Frances Clark^2^, Billy T. George^2^, Lauren Gregory^2^, David Tess^2^, James R. Gosset^3^, Jennifer Liras^3^, Emily Geishecker^1^, R. Scott Obach^2^, Murat Cirit^1^@

* Co-first authors

@ corresponding author

Mailing address: 299 Washington Street Woburn, MA 01801

^1^Javelin Biotech Inc., 299 Washington street, Woburn, Massachusetts 01801, United States

^2^Pfizer Global Research and Development, Groton Laboratories, Eastern Point Road, Groton, Connecticut 06340, United States

^3^Pfizer Worldwide Research and Development, 610 Main Street, Cambridge, Massachusetts 02139, United States

***Supplementary Materials & Methods***

**Liver Tissue Chip Culture:** Prior to use, the chips were pre-surface treated and sterilized using ethanol or gamma irradiation. On the day before seeding, the cell culture chambers were coated with ECM solution, containing 100μg/mL of rat tail collagen I (Corning) and 25 µg/mL fibronectin (Sigma Aldrich) and incubated at 4ᵒC overnight followed by at least 2-hour incubation at 37ᵒC prior to seeding. Cryopreserved Primary Human Hepatocytes (PHH, BioIVT) were thawed in Cryopreserved Hepatocyte Recovery Medium (CHRM, Gibco) and centrifuged at 100g for 8 min in pre-warmed CHRM. The cells were re-suspended at a density of 0.86 e6/mL cells in appropriate volume of pre-warmed Hepatocyte Plating Medium (HPM), that included Williams’ E Medium (WEM, Gibco) and Primary Hepatocyte Thawing and Plating Supplements (Gibco), and directly seeded to the cell culture chamber (215K cells/chip).

The hepatocytes were incubated undisturbed for 4hrs allowing cell attachment, after which the culture was washed with cold Hepatocyte Maintenance Medium (HMM), prepared by adding Hepatocyte Maintenance Supplements (Gibco) to WEM, and overlayed with 0.35 μg/mL Matrigel (Corning) prepared in cold HMM after which the chips were incubated overnight at 37ᵒC. Next day, the culture chamber was sealed with chamber lid, and 1.7mL of HMM was added to the chip using a syringe through the fill port. The chips were then connected to the support controller and set to continuous flow at 2mL/hr. The chips were maintained for long-term culture by replacing 400μL spent media every 2-3 days and monitored using brightfield microscopy.

**Phenotypic Characterization:** Albumin and urea production in the LTC was measured by monitoring the levels in culture medium collected from the chip during partial media changes. For albumin quantification, the media samples were diluted at 1:250 and quantified according to vendor protocols for the R-PLEX Human Albumin Antibody Set using the MESO QuickPlex SQ120MM plate reader (Meso Scale Discovery). Urea concentrations of each media sample were measured using the QuantiChrom Urea Assay Kit (BioAssay Systems). Vendor recommendations for low urea samples were used and optical density was measured using a SpectraMax M3 Multi-mode Spectrophotometer at 430nm. Albumin and urea concentrations in media were corrected for number of cells, chip volume, and partial media change volume and frequency to yield daily albumin and urea production rates per million cells.

Several major isoforms of Cytochrome P450 enzyme activity were determined using a probe substrate cocktail of tacrine (CYP1A2; 3 μM), diclofenac (CYP2C9; 90 μM), omeprazole (CYP2C19; 3 μM), dextromethorphan (CYP2D6; 20 μM), and midazolam (CYP3A4; 3 μM), (Table S1). On days 3, 8, and 13 of culture, the spent medium in the LTC was completely replaced with the above cocktail media and incubated for 3 hours under flow at 37ᵒC. At the end of incubation, media samples were collected, cocktail media was replaced with fresh maintenance media (HMM), and the LTC was connected back to flow. Metabolism was assessed by quantification of phase 1 metabolites (1’-hydroxytacrine, 4’-hydroxydiclofenac, 5’-hydroxyomeprazole, dextrorphan, and 1’-hydrozymidazolam, respectively) by LC-MS/MS.

**Table S1. Summary of the probe substrates and concentration used in the CYP probe substrate assay to determine the activity levels of CYP isoform.**

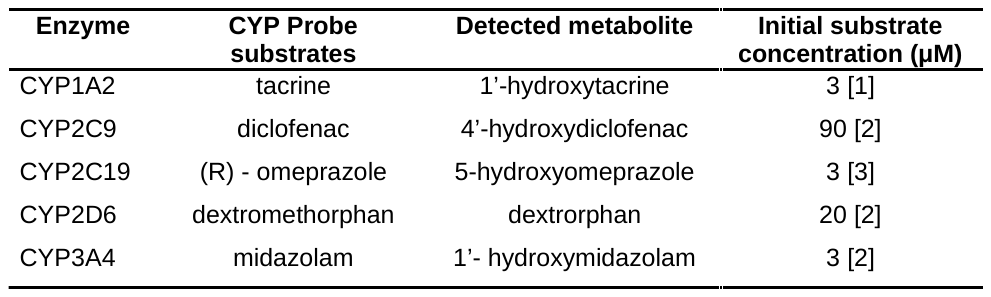

To visualize bile canaliculi in hepatocyte culture, live staining was performed by incubating the culture with 5-chloromethylfluorescein diacetate (CMFDA, Invitrogen). The spent media in the LTC was replaced with CMFDA containing media by diluting 10mM stock solution 1:500 in maintenance media. The chips were incubated under flow for 2-hours, and the CMFDA media was replaced with fresh HMM prior to imaging. The chips were imaged directly using a fluorescent microscope with 488nm fluorescence. Brightfield images were taken simultaneously to overlay the stained Bile canaliculi images.

For IHC staining, the cultured liver tissue on MPS was fixed with 4% paraformaldehyde for 15 min at room temperature (RT) or overnight at 4ᵒC and permeabilized by incubation in 0.2% Triton-X for 10 min at RT. Non-specific antibody binding was blocked by incubation in Protein Block Solution (Abcam) for 30 min in a rocker/shaker at RT. The primary antibodies diluted in antibody diluent (Abcam) were added and incubated overnight at 4ᵒC. The hepatocytes were stained with E-Cadherin (Abcam), F-Actin (Phalloidin, Cayman Chemical Company), CK-18 (Abcam), MRP-2 (Fisher Scientific), and CYP3A4 (ThermoFisher). Following primary incubation, the tissues were washed in PBS and incubated for 1 hr with anti-rabbit, anti-mouse, Alexa Fluor 488, 594 and 647 secondary antibodies (Invitrogen) as appropriate in antibody diluent (1:200 dilution). Cells were counterstained with DAPI for 7 min at RT and washed with PBS prior to fluorescent imaging. The tissue in the LTC was imaged using a Zeiss LSM880 upright confocal microscope.

**Quantitative real-time PCR (qPCR):** Total RNA was isolated from cells in the liver MPS using Invitrogen PureLink RNA Mini kit (Invitrogen) following the vendor's recommended protocol. Briefly, the hepatocyte monolayer was lysed using a lysis buffer and the lysate was collected from the chip and processed. RNA was converted to cDNA using Applied Biosystems High-Capacity cDNA Reverse Transcription Kit (Applied Biosystems) which was then loaded into a custom Taqman Array plate (ThermoFisher) following the vendor's recommended protocol. The cDNA was analyzed using StepOnePlus Real-Time PCR System (ThermoFisher). Threshold Ct values for each sample were set at 35 for the analysis. RPLO and 18S were selected as house-keeping genes and their expression levels were calculated in parallel with the genes of interest.

**Drug preparation:** Drug stock solutions were prepared by solubilizing powdered drug in DMSO in a concentration at least 1000-times higher than the required final concentration and stored at -80ᵒC until use. Midazolam and alprazolam (Sigma-Aldrich) were pre-solubilized from vendor in methanol and were mixed with DMSO to obtain a stock solution of 1mM concentration on the day of drug addition. Drug cocktails of two or more drugs were prepared by reconstituting the stock solution of each drug in media to reach a final concentration of 1 µM with less than 0.1%DMSO in the media, except propranolol which was reconstituted to 0.1 µM. Drugs in the cocktails were chosen in specific combinations to avoid any effect or risk of Drug-Drug interactions (DDI) that could affect the clearance prediction (Table S2). For example, drugs metabolized by the same CYP isoform were not in the same cocktail neither were the drugs that are substrates to the same transporter. We also confirmed analytical (LC-MS/MS) compatibility to be quantified in the same mixture.

**Table S2.** **List of drug cocktails used in the LTC drug study with clearance characteristics of the drug for hepatic tissue.**

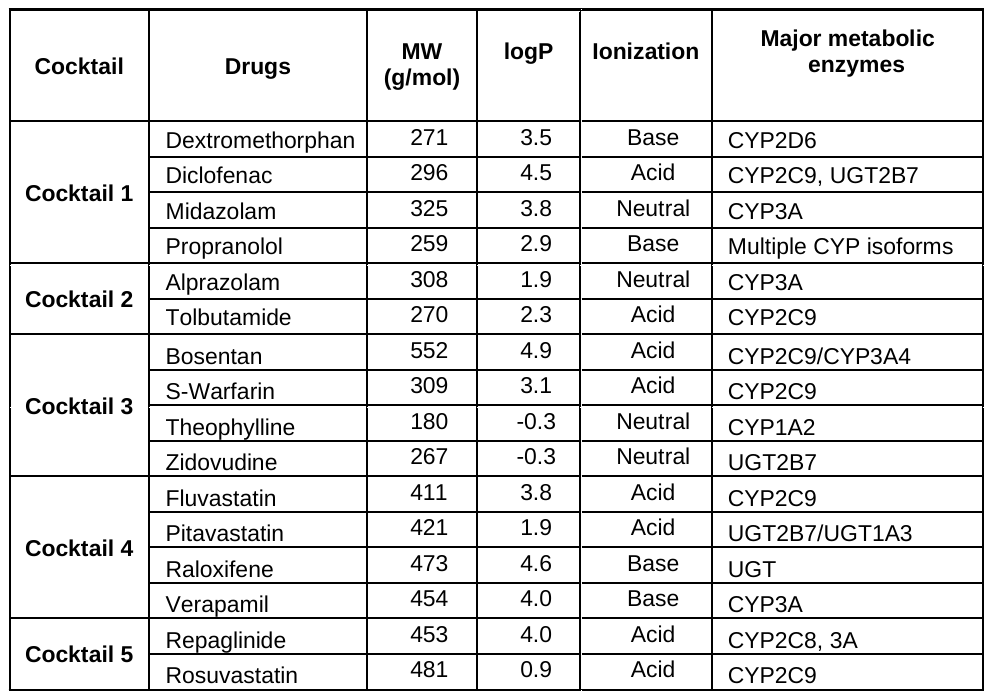

**Bioanalysis:** HPLC grade water, HPLC grade water containing 0.1% (v:v) formic acid HPLC grade acetonitrile (ACN), and HPLC grade ACN containing 0.1% (v:v) formic acid were obtained from Fisher Scientific. The following reference material: midazolam, diclofenac, tolbutamide, propranolol, dextromethorphan, alprazolam, rosuvastatin, theophylline, alprazolam, S-warfarin, zidovudine, dextromethorphan, pitavastatin, bosentan, fluvastatin, verapamil, raloxifene, and repaglinide were either provided by Pfizer, Inc. or purchased from Caymen Chemical, Michigan, United States. Stock solutions of each compound were first prepared in DMSO in 1 to 90 mM range. 100 μM cocktailed stocks were prepared in 50:50 (v:v) ACN:DMSO for analyte stock solutions < 1mM. All stock solutions that were ≥ 10 mM were combined to create a 1 mM intermediate solution in 50:50 (v:v) ACN:DMSO. Standard calibration curves were prepared by serial dilution in HMM over the range of 0.1 to 5000 nM. Standards were included in the calibration curve if they had a %RE of ≤ 20%.

Study samples were prepared by diluting 5 to 10 µL of HMM sample with 100 µL of acetonitrile containing internal standards and the sample aliquot volume was determined by required sensitivity. Refer to Table S4 - S7 for internal standard compounds. The mixture was centrifuged at 3000 RPM for 5 minutes. 50 µL of the supernatant was diluted with water containing 0.1% (v:v) formic acid, mixed, and then injected onto the LC-MS/MS system. The chromatographic system was allowed to equilibrate with each initial mobile phase condition prior to analysis.

The chromatographic system included a Shimadzu Nexera X2 CBM-20A communication bus module, Shimadzu Nexera X2 LC-30AD pumps, Shimadzu Nexera X2 SIL-30ACMP autosampler, and Shimadzu DGU-20A5 degassers. Mobile phase A consisted of water containing 0.1% (v:v) formic acid and mobile phase B consisted of acetonitrile containing 0.1% (v:v) formic acid. The flow rate and column temperatures were 0.5 ml/min and 40°C, respectively. The chromatographic separations were achieved for Study A using a Waters Acquity UPLC HSS T3 column, 1.8 µM, 2.1x50mm, P/N 186003538. A Waters Acquity BEH C18 column (2.1 mm x 50 mm, 1.7 µm, P/N 186002350 was employed for studies B, C, D and E. The gradient programs are tabulated in Table S3.

The mass spectrometric detector utilized to conduct sample analysis for studies A and B was a Sciex Triple Quad™ 5500 equipped with a IonDrive™ Turbo V source. The source conditions were as follows: positive ion electrospray, CUR 20, CAD 7, IS 3000, TEM 600, GS1 50, GS2 60, EP 10, and CXP 14 for studies A and B. For study E, the source conditions were as follows: positive ion electrospray, CUR 20, CAD 9, IS 5500, TEM 550, GS1 50, GS2 60, EP 10, and CXP 14. A Sciex Triple Quad™ 6500 Plus equipped with a IonDrive™ Turbo V source was utilized for studies C and D. The source conditions were: positive ion electrospray, CUR 20.0, CAD 8, IS 4500, TEM 600, GS1 50, GS2 60, EP 10, and CXP 10.

The data acquisition was performed using multiple reaction monitoring (MRM). MRM transitions can be found in Tables S4 to S7. Data acquisition and chromatographic review were performed using Applied Biosystems/MDS Sciex Analyst, version 1.7.2. The calibration curves utilized 1/x^2^ weighting.

**Table S3 Chromatographic Gradients**

**Studies A and B**

| **Time (min)** | **Module** | **Event** | **Parameter** |
| --- | --- | --- | --- |
| 0.01 | Pumps | Pump B Conc. | 5 |
| 0.30 | Pumps | Pump B Conc. | 5 |
| 2.30 | Pumps | Pump B Conc. | 95 |
| 2.60 | Pumps | Pump B Conc. | 95 |
| 2.70 | Pumps | Pump B Conc. | 5 |
| 3.00 | Controller | Stop |  |

**Study C and D**

| **Time (min)** | **Module** | **Event** | **Parameter** |
| --- | --- | --- | --- |
| 0.01 | Pumps | Pump B Conc. | 2 |
| 0.70 | Pumps | Pump B Conc. | 2 |
| 2.10 | Pumps | Pump B Conc. | 95 |
| 2.60 | Pumps | Pump B Conc. | 95 |
| 2.70 | Pumps | Pump B Conc. | 2 |
| 4.00 | Controller | Stop |  |

**Study E**

| **Time (min)** | **Module** | **Event** | **Parameter** |
| --- | --- | --- | --- |
| 0.01 | Pumps | Pump B Conc. | 5 |
| 0.30 | Pumps | Pump B Conc. | 5 |
| 3.30 | Pumps | Pump B Conc. | 95 |
| 3.60 | Pumps | Pump B Conc. | 95 |
| 3.70 | Pumps | Pump B Conc. | 5 |
| 4.00 | Controller | Stop |  |

**Table S4. Study A** **MRM Parameters**

| **MRM Parameters** | | | | | |
| --- | --- | --- | --- | --- | --- |
| **Q1** | **Q3** | **Dwell time** | **Compound** | **DP** | **CE** |
| **Analytes** | | | | | |
| 411.900 | 224.000 | 30.0 | Fluvastatin^1^ | 15.000 | 30.000 |
| 421.900 | 290.100 | 30.0 | Pitavastatin^2^ | 60.000 | 42.000 |
| 474.300 | 112.100 | 30.0 | Raloxifene^3^ | 122.000 | 44.000 |
| 455.500 | 165.400 | 30.0 | Verapamil^2^ | 60.000 | 45.000 |
| 453.200 | 162.400 | 30.0 | Repaglinide^2^ | 55.000 | 30.000 |
| 482.100 | 258.100 | 30.0 | Rosuvastatin^2^ | 140.000 | 40.000 |
| 553.000 | 202.000 | 30.0 | Bosentan | 58.000 | 42.000 |
| 309.10 | 251.100 | 30.0 | S-Warfarin^2^ | 20.000 | 25.000 |
| 180.600 | 103.000 | 30.0 | Theophylline-3^4^ | 20.000 | 15.000 |
| 268.100 | 127.000 | 30.0 | Zidovudine^3^ | 20.000 | 19.000 |
| **Internal Standards** | | | | | |
| 358.300 | 139.200 | 30.0 | Indomethacin | 46.000 | 26.000 |
| 472.100 | 436.000 | 30.0 | Terfenadine | 80.000 | 30.000 |
| 268.100 | 116.100 | 30.0 | Metoprolol | 60.000 | 25.000 |
| 288.200 | 243.200 | 30.0 | Zolmitriptan | 65.000 | 25.000 |
| 419.200 | 285.200 | 30.0 | Simvastatin | 40.000 | 15.000 |

^1^Internal standard used was indomethacin.

^2^Internal standard used was terfenadine.

^3^Internal standard used was metoprolol.

^4^Internal standard used was zolmitriotan.

**Table S5. Study B MRM Parameters**

| **MRM Parameters** | | | | | |
| --- | --- | --- | --- | --- | --- |
| **Q1** | **Q3** | **Dwell time** | **Compound** | **DP** | **CE** |
| **Analytes** | | | | | |
| 553 | 202 | 30.0 | Bosentan^1^ | 58.0 | 42.0 |
| 268.08 | 126.96 | 30.0 | Zidovudine^2^ | 20.0 | 19.0 |
| 181.1 | 124.2 | 30.0 | Theophylline-2/-3^3^ | 20.0 | 15.0 |
| 309.1 | 251.1 | 30.0 | S Warfarin^4^ | 20.0 | 25.0 |
| **Internal Standards** | | | | | |
| 358.3 | 139.2 | 15.0 | Indomethacin | 46.0 | 26.0 |
| 268.1 | 116.1 | 15.0 | Metoprolol | 60.0 | 25.0 |
| 288.2 | 243.2 | 15.0 | Zolmitriptan | 65.0 | 25.0 |
| 472.1 | 436 | 15.0 | Terfenadine | 80.0 | 30.0 |

^1^Internal standard used was indomethacin.

^2^Internal standard used was metoprolol.

^3^Internal standard used was zolmitriptan.

^4^Internal standard used was terfenadine.

**Supplementary Table 6. Studies C and D MRM Parameters**

| **MRM Parameters** | | | | | |
| --- | --- | --- | --- | --- | --- |
| **Q1** | **Q3** | **Dwell time** | **Compound** | **DP** | **CE** |
| **Analytes** | | | | | |
| 272.3 | 215.2 | 15.0 | Dextramethorphan^1^ | 80.0 | 35.0 |
| 326.1 | 222.2 | 15.0 | Midazolam^1^ | 60.0 | 60.0 |
| 260.2 | 116.2 | 15.0 | Propranolol^1^ | 80.0 | 30.0 |
| 296.1 | 214.0 | 15.0 | Diclofenac^1^ | 70.0 | 40.0 |
| 309.2 | 205.2 | 15.0 | Alprazolam^2^ | 70.0 | 60.0 |
| 271.0 | 172.0 | 15.0 | Tolbutamide^2^ | 66.0 | 18.0 |
| **Internal Standards** | | | | | |
| 268.1 | 116.1 | 15.0 | Metoprolol | 60.0 | 25.0 |
| 472.1 | 436.0 | 15.0 | Terfenadine | 80.0 | 30.0 |

^1^Internal standard used was metoprolol.

^2^Internal standard used was terfenadine.

**Table S7. Study E MRM Parameters**

| **MRM Parameters** | | | | | |
| --- | --- | --- | --- | --- | --- |
| **Q1** | **Q3** | **Dwell time** | **Compound** | **DP** | **CE** |
| **Analytes** | | | | | |
| 296.1 | 214.0 | 30.0 | Diclofenac^1^ | 70 | 40 |
| 312.0 | 266.0 | 30.0 | 4-Hydroxy^2^ Diclofenac | 50 | 20 |
| 272.3 | 215.2 | 30.0 | Dextromethorphan^3^ | 80 | 35 |
| 258.1 | 201.1 | 30.0 | Dextrorphan^4^ | 40 | 31 |
| 326.0 | 291.0 | 30.0 | Midazolam^3^ | 136 | 39 |
| 342.0 | 325.0 | 30.0 | 4-Hydroxy Midazolam^3^ | 106 | 31 |
| 199.1 | 171.2 | 30.0 | Tacrine^4^ | 50 | 30 |
| 215.0 | 197.0 | 30.0 | Hydroxy Tacrine^5^ | 50 | 30 |
| 823.1 | 791.3 | 30.0 | Rifampicin^2^ | 85 | 31 |
| **Internal Standards** | | | | | |
| 358.3 | 139.2 | 30.0 | Indomethacin | 46 | 26 |
| 268.1 | 116.1 | 30.0 | Metoprolol | 60 | 25 |
| 260.2 | 116.2 | 30.0 | Propranolol | 80 | 30 |
| 472.1 | 436.0 | 30.0 | Terfenadine | 80 | 30 |
| 288.2 | 243.2 | 30.0 | Zolmitriptan | 65 | 25 |

^1^Internal standard used was Indomethacin.

^2^Internal standard used was Terfenadine.

^3^Internal standard used was Propranolol.

^4^Internal standard used was Metoprolol.

^5^Internal standard used was Zolmitriptan.

**Statistical Analysis:** Each study arm in each experiment contained at least four LTCs as biological replicates. Chip-to-chip variability was assessed by calculating the coefficient of variation (CV%) for each biochemical assay (albumin, and urea) and predicted intrinsic clearance values (${CL}_{int}$) for tested drugs. Statistical significance between drug study window experiments was tested using one-way ANOVA non-parametric testing, and the study arms for parametric data were tested using Student’s t-test or one-way. Aggregate statistics of the independently assessed drug depletion data from each chip were then computed across chips for each drug using geometric mean and CV [4]. The fold changes in gene expression using RT-PCR were analyzed via the 2−△△CT method normalizing the expression to house-keeping genes; the change in expression for the genes of interest was calculated in comparison to LTC expression on day 3.

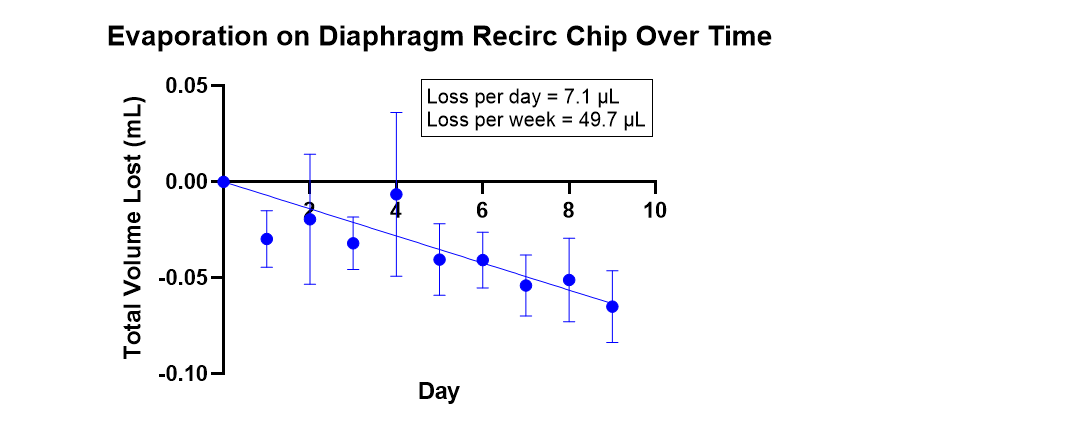

**Figure S1. Rate of evaporation measured on LTC over 9 days showing insignificant volume changes over time due to evaporation.**

**RT-PCR characterization of pharmacokinetic genes.**

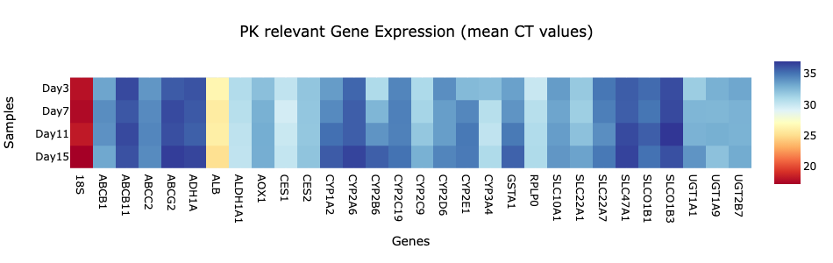

**Figure S2. RT-PCR characterization of PK relevant genes expressing stable Mean CT values on LTC.** Stable expressions (lower the CT, higher the expression) can be seen across the gene panel for days 3, 7, 11, and 15. Housekeeping genes 18S and RPLP0 are used to normalize the expression for all the PK relevant genes.

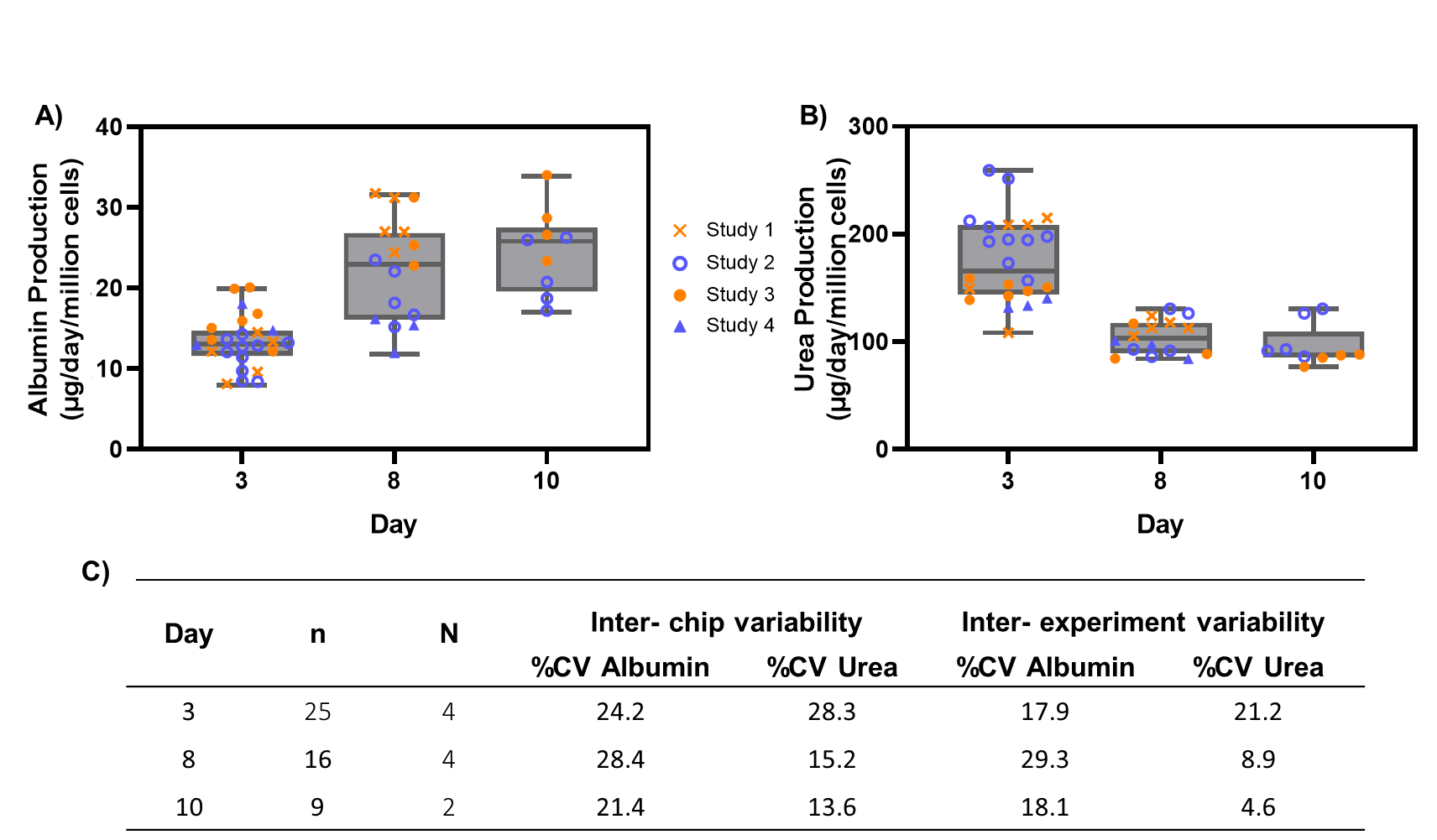

**Figure S3. Reproducibility and repeatability of liver tissue chips validated with functional markers.** (A) Albumin and (B) Urea production values of multiple chips run within the same experiment or different experiment group and C) Table to summarize the coefficient of variation (CV%) between functional markers of different LTC over 10 days showing <28.4% for albumin and <28.3% for urea production for inter-chip variability and <29.3% for albumin and <21.2% for urea production for inter-experiment variability over four individual experiments.

**Modeling the effect of recirculation on drug metabolism studies.**

*Flow-through chip configuration:*

The change in drug concentration in the flow-through chip was modeled using one compartment for the cell chamber (with concentration $C_{cells}$ [μM] and volume $V_{cells}$ [mV]) and one for output media collection (with concentration $C_{out}$ [μM] and volume $V_{out}$ [mV]) according to the equations:

$$\frac{dC_{cells}}{dt}=\frac{1}{V_{cells}}\left( Q_{sys}*\left( C_{0}-C_{cells} \right) \right)-SF*{CL}_{intu}*{fu}_{media}*C_{cells}$$

$$\frac{dC_{out}}{dt}=\frac{1}{V_{out}}\left( Q_{sys}*\left( C_{cells}-C_{out} \right) \right)$$

where

SF = $N_{cells}$*(60 min/h)*(1000 μl/ml)

Initial conditions: $C_{cells}\left( t=0 \right)=C_{o}=1\mu M$, $C_{out}\left( t=0 \right)=0$

Units: ${CL}_{intu}$ [μl/min/MC], C [μM], t [h], V [mL]

Drug was assumed to enter continuously with concentration $C_{0}$, flow unidirectionally through the chip with systemic flow rate $Q_{sys}$ into the media collection reservoir, and to be metabolized by the hepatocytes in the cell chamber with intrinsic metabolic clearance ${CL}_{intu}$. The kinetic profile was plotted over 6 hours to show the transient and steady state behavior of the system for drugs with different rates of metabolism.

*Recirculation chip configuration:*

The chip with recirculation was modeled using two compartments for the cell chamber (with concentration $C_{cells}$ [μM] and volume $V_{cells}$ [mV]) and one representing the recirculation path (with concentration $C_{recirc}$ [μM] and volume $V_{recirc}$ [mV]) according to the equations:

$$\frac{dC_{cells}}{dt}=\frac{1}{V_{cells}}\left( Q_{sys}*\left( C_{recirc}-C_{cells} \right) \right)-SF*{CL}_{intu}*{fu}_{media}*C_{cells}$$

$$\frac{dC_{recirc}}{dt}=\frac{1}{V_{recirc}}\left( Q_{sys}*\left( C_{cells}-C_{recirc} \right) \right)$$

Initial conditions: $C_{cells}\left( t=0 \right)=C_{o}=1\mu M$, $C_{out}\left( t=0 \right)=1\mu M$

Drug was assumed to be dosed initially with concentration $C_{0}$, flow through the chip in a closed loop with systemic flow rate $Q_{sys}$, and to be metabolized by hepatocytes in the cell chamber. The kinetic profile was plotted over 8 days to show the drug depletion that is observable with the recirculation design.

*Chip parameters used for both models:*

$N_{cells}=$ 215,000

$Q_{sys}$ = 2 mL/h

$V_{cells}=$ 0.4 mL

$V_{out}=V_{recirc}=$ 1.3 mL

*Drug-related parameters used for both models:*

${fu}_{media}=$ 1

${CL}_{intu}$ = 0.08, 5, or 30 μl/min/million cells (MC)

Intrinsic clearance values were chosen to model metabolism for compounds with fast clearance (30 μl/min/MC), moderate clearance (5 μl/min/MC), and slow clearance producing 10% depletion over 8 days (0.08 μl/min/MC).

Simulations were performed using the DynaSim toolbox [5] in MATLAB R2019b.

**Prediction of in vivo hepatic clearance.**

Drug-related parameters extracted from MPS studies can be scaled to predict clinical parameters using in vitro-in vivo translation [6]. The typical value of unbound intrinsic clearance ($\underline{CL}_{int(u)}$) determined for each drug from the pharmacokinetic analysis of the individual-donor in vitro data was scaled up to a human liver equivalent unbound intrinsic clearance (${CL}_{int(u),H}$) using

$${CL}_{int(u),H}=\underline{CL}_{int(u)}\cdot HC\cdot LW$$

where HC is the human hepatocellularity of 120 million cells / g of liver [7] and LW is the average human liver weight of 25.7g / kg of body weight [8]. The hepatic clearance (referring to whole blood concentrations) was then predicted (${CL}_{H(pred)}$) using the Parallel Tube (PT) liver model [9]; for completeness, the hepatic clearance was also calculated using the Well-Stirred (WS) model:

$${CL}_{H(pred,PT)}=Q_{H}\cdot\left( 1-e^{-{\cdot fu}_{b}\cdot{CL}_{int\left( u \right),H}/Q_{H}} \right)$$

$${CL}_{H(pred,WS)}=\frac{Q_{H}{\cdot fu}_{b}\cdot{CL}_{int\left( u \right),H}}{Q_{H}{+fu}_{b}\cdot{CL}_{int\left( u \right),H}}$$

where is the average hepatic blood flow of 20.7 mL/min/kg of body weight [10] and ${fu}_{b}$ is the fraction of drug which is unbound in blood. The fraction unbound in blood (${fu}_{b}$) was calculated for each compound from the known fraction unbound in the plasma (${fu}_{p}$) and blood-to-plasma ratio ($Rbp$) (Table S8) according to the equation ${fu}_{b}={fu}_{p}/Rbp$. The predicted hepatic clearance (${CL}_{H(pred)}$) values were then compared to observed hepatic clearance (${CL}_{H(obs)}$) values (referring to whole blood concentrations), which were calculated under the assumption of no extra-hepatic metabolism

$${CL}_{H(obs)}={CL}_{T(obs)}-{CL}_{R(obs)}$$

where ${CL}_{T(obs)}$ is the total clearance (referring to whole blood) that has been observed in humans (see values in Table S9) and ${CL}_{R(obs)}$ is the renal clearance determined from drug excreted unchanged in urine (see values in Table S8). Clinical intrinsic clearance estimates (${CL}_{int(obs)}$) were further derived for 10 compounds that were not flow rate limited by applying the inverse parallel tube model to the clinical data (${CL}_{H(obs)}$). The overall agreement between the observed and the predicted hepatic clearance values was determined by the calculation of the average absolute fold error (AAFE) across all the evaluated compounds (N=10 or 12) using

$$AAFE={10}^{\frac{1}{N}\sum_{q=1}^{N} \left| log\frac{{CL}_{H\left( obs \right),q}}{{CL}_{H\left( pred \right),q}} \right|}$$

$$AAFE={10}^{\frac{1}{N}\sum_{q=1}^{N} \left| log\frac{{CL}_{int\left( obs \right),q}}{{CL}_{int\left( pred \right),q}} \right|}$$

**Table S8. Experimental and literature values used for scaling in vitro parameters.** fu_media_, fu_p_, and Rbp were experimentally determined. ${CL}_{T}$ and ${CL}_{R}$ values were obtained from the following references: [11] Smith 1984; [12] Moghadamnia 2003; [13] Lombardo 2018; [14] Hatorp 1998; [15] Chen 2020; [16-17] FDA; [18] Scotcher 2016.

| Drug | fu_media_ | fu_p_ | Rbp | CL_T_ | CL_R_ |
| --- | --- | --- | --- | --- | --- |
| Alprazolam | 0.7 | 0.35 | 0.87 | 0.71^11^ | 0.1^15^ |
| Dextromethorphan | 0.792 | 0.455 | 1.2 | 18.4^12^ | 0.006^15^ |
| Diclofenac | 0.0387 | 0.004 | 0.68 | 3.5^13^ | 0^16^ |
| Midazolam | 0.488 | 0.053 | 0.55 | 5.3^13^ | 0^15^ |
| Propranolol | 0.98 | 0.297 | 0.754 | 12^13^ | 1.2^15^ |
| Raloxifene | 0.0886 | 0.017 | 0.78 | 10.8^13^ | 1.47^15^ |
| Repaglinide | 0.0224 | 0.015 | 0.74 | 7.39^14^ | 0^17^ |
| S-Warfarin | 0.498 | 0.013 | 0.595 | 0.055^13^ | 0^15^ |
| Theophylline | 0.608 | 0.654 | 0.73 | 0.86^13^ | 0.0 86^15^ |
| Tolbutamide | 0.6 | 0.026 | 0.562 | 0.21^13^ | 0^15^ |
| Verapamil | 0.35 | 0.186 | 0.665 | 18^13^ | 0.36^18^ |
| Zidovudine | 0.875 | 0.883 | 1 | 25^13^ | 5.67^15^ |

**Table S9**. Results of the in vitro to in vivo comparison.

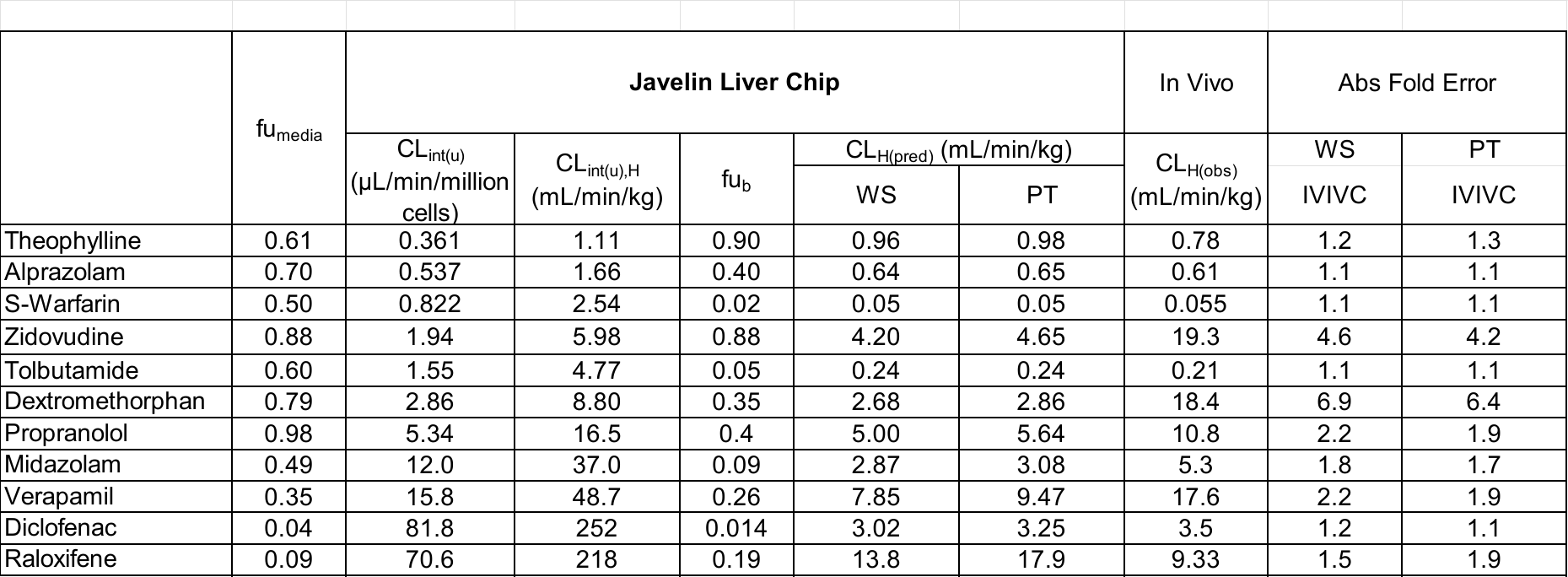

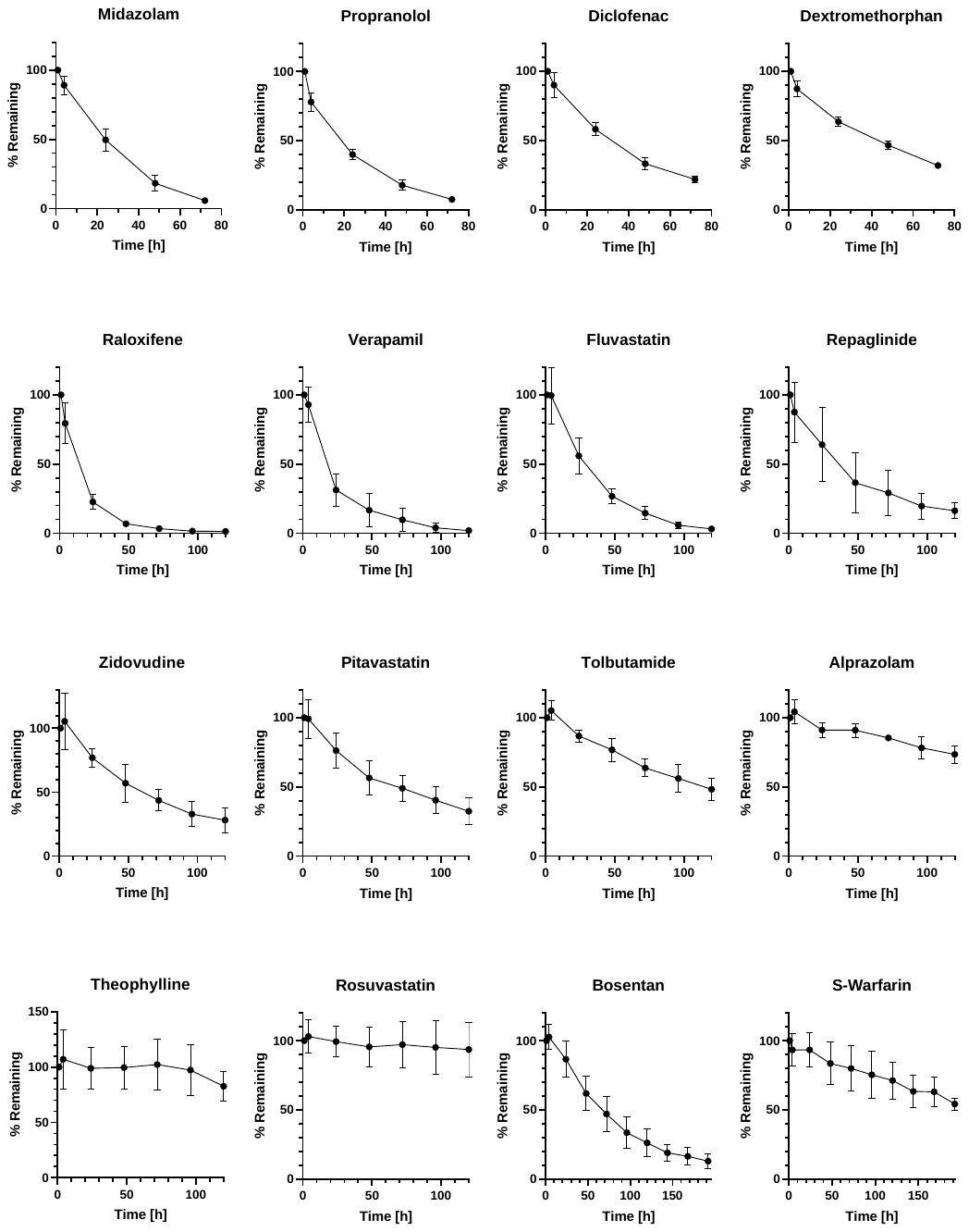

**Figure S4. Kinetic profiles from metabolism studies used for in vivo prediction.**
